# AMMPER-2: A spatially explicit agent-based model of microbial radiobiology with redox dye simulation

**DOI:** 10.64898/2026.10.08.757403

**Authors:** Daniel Palacios, Pramesh Sharma, Madeline Marouš, Danny Mejia, Lauren C. Liddell, Diana M. Gentry, Sergio R. Santa Maria, Jessica A. Lee

**Affiliations:** Quantitative and Computational Biosciences, Baylor College of Medicine, Houston, Texas 77030, USA; Department of Pediatrics, Baylor College of Medicine, Houston, Texas 77030, USA; Jan and Dan Duncan Neurological Research Institute, Texas Children’s Hospital, Houston, Texas 77030, USA; Department of Computer Science, University of California Davis, Davis, CA, USA; Department of Information Sciences and Technology, Penn State Berks, Reading, PA, USA; Miami Dade College, Miami, FL, USA; Logyx LLC, Mountain View, CA, USA; Earth Sciences Division, NASA Ames Research Center, Moffett Field, CA, USA; Space Biosciences Division, NASA Ames Research Center, Moffett Field, CA, USA

**Author notes:** Address correspondence to Jessica Lee. Daniel Palacios and Pramesh Sharma contributed equally to this work. Author order was determined by D.P. and the senior author, with concurrence by all other authors.

## Abstract

To reduce health risks for human space exploration, it is important to model the effects of deep-space radiation on biological systems. The budding yeast *Saccharomyces cerevisiae* is a common model organism in space radiobiology, but little is known about how radiation damage to individual cells translates into the population-level effects that experiments measure. The Agent-Based Model for Microbial Populations Exposed to Radiation (AMMPER) is a spatially explicit agent-based model that simulates the effects of ionizing radiation on *S. cerevisiae* cells and produces population-level results. Its goal is to improve experiment design and data interpretation. AMMPER-1 demonstrated that agent-based simulations can qualitatively reproduce the effects of proton radiation on the growth rate of wild-type and DNA-repair mutant yeast. Here, we present AMMPER-2, which addresses limitations of the first model and expands its capabilities. AMMPER-2 incorporates the dynamics of the redox dye alamarBlue, a data type commonly generated in spaceflight microbiology experiments, and models the fate of reactive oxygen species in more detail. It also adds an improved graphical interface and runs faster. AMMPER-2 remains easy to execute in Python on a standard personal laptop, providing space biologists a tool to explore experimental conditions, generate hypotheses, and interpret data from microbial space radiation experiments.

**IMPORTANCE:** Understanding how deep-space radiation affects living systems is essential for safe long-duration human space exploration. However, the link between radiation damage at the level of the individual cell and the population-level growth and metabolic signals measured in spaceflight experiments remains poorly understood. Computational tools that connect these scales can improve experiment design and data interpretation. This manuscript describes an update to AMMPER, a previously published computational modeling tool. The intended users are scientists, educators, and students interested in the biological effects of radiation on organisms, and the tool prioritizes ease of use and accessibility over complexity and precision. This model is a work in progress, and the manuscript focuses on new features and lessons learned rather than scientific findings.

## INTRODUCTION

### Microbial experiments in deep space

Microorganisms feature prominently in experimental space biology [1, 2, 3]. Their small size, rapid generation time, genetic manipulability and tolerance of stasis make them convenient models for the response to spaceflight stressors such as radiation and altered gravity. Microbiomes are an unavoidable part of the habitable environments occupied by astronauts, so studying microbial behavior in space is necessary for establishing healthy habitats [4, 5, 6, 7, 8]. Microorganisms will also carry out essential functions in long-distance travel, including bioregenerative life support, food production, biomining, and biosynthesis of products such as pharmaceuticals [9, 10, 11, 12, 13, 14]. Those functions require empirical testing in space.

A platform particularly well suited to microorganisms is the CubeSat, a small modular satellite. Experimental progress in space microbiology and astrobiology is limited in part by access to space aboard human-tended spacecraft and space stations, a limitation CubeSats help address [15]. A CubeSat payload can carry out a biological experiment independently of astronaut support [16]. Examples include GeneSat-1 [17], PharmaSat [18], O/OREOS [19], EcAMSat [20], SporeSat [21] and BioSentinel [16]; the platform and its microbiology applications are reviewed elsewhere [22, 15]. These experiments are automated, self-contained, small in volume and often require no sample return. They form the basis for the first experiments on the distinctive radiation environment beyond Low Earth Orbit (LEO), where opportunities for crew-based experimentation are extremely limited. BioSentinel was launched as a secondary payload on *Artemis I* into a trailing heliocentric orbit and is, at the time of writing, more than 77 million km from Earth and receding. LEIA (the Lunar Explorer Instrument for space biology Applications) is being developed with very similar instrumentation for delivery to the lunar surface by an uncrewed lander, NET 2027 [23, 24].

Both missions use the BioSensor, a microfluidic culturing device accommodating at least 256 samples of *∼*100 *µ*L. Yeast cells are stored desiccated in individual wells of fluidic cards, in stasis for launch and transit. The experiment is initiated by pumping culture medium into the wells, after which growth and metabolism are monitored by optical absorbance at three wavelengths per well at frequent timepoints. BioSentinel and half of the LEIA samples, like several prior spaceflight experiments, include the redox-sensitive dye resazurin (trade name alamarBlue) in the medium. Measurements at 570 nm and 630 nm track the dye’s color change from blue to pink to colorless as it is reduced by electron carriers produced through cellular metabolism [25]. Optical density is read at 850 nm to avoid interference. These missions are automated and data-rich. Interpreting how cellular radiation damage gives rise to the population-level metabolic and growth signals they measure is difficult, and that difficulty motivates the development of computational tools such as AMMPER.

### The space environment: deep space ionizing radiation

Of the many challenges the space environment imposes on biological systems, ionizing radiation is of particular concern, because its effects are universal across the tree of life and effective mitigation strategies are lacking [26, 3]. The environment beyond LEO differs from the LEO environment in which most space biology experimentation has been conducted to date and poses greater health risks. This holds on the Moon, on Mars, and in transit to those destinations. Its characteristic component is Galactic Cosmic Radiation (GCR), formed outside the solar system by extreme events such as supernovae and by stellar winds. GCR arrives from all directions and is composed principally of protons, followed by *α* particles, with 1% heavier nuclei [27]. It causes direct DNA damage including double- and single-strand breaks [28, 29]. It also acts indirectly, interacting with the water in cell culture medium to create free radicals or Reactive Oxygen Species (ROS) such as *OH* and *H*_2_*O*_2_. These impose oxidative stress and at high concentrations induce apoptosis [28, 30, 31].

### AMMPER-1

The Agent-Based Model for Microbial Populations Exposed to Radiation (AMMPER) was developed to help researchers and educators understand the biological effects of ionizing radiation in microbial cells [32]. Written in Python, it treats *S. cerevisiae* cells as individual agents. The model combines DNA damage, a DNA repair probability model, Brownian motion, cell replication, ROS modeling and three radiation environment models (Deep Space Proton, NSRL GCRSim Proton, and 150 MeV Proton). From these it predicts cell health outcomes and population-level dynamics, including growth curves and growth rates. Spatially dependent energy deposition events are generated with the Relativistic Ion Tracks (RITRACKS) model [33, 34]; these liberate ROS molecules and damage nearby cells both directly and indirectly (Fig. **1**A). Exposure occurs either at a specified generation (the 150 MeV proton and GCRSim scenarios) or chronically at every generation (Deep Space Proton). AMMPER models the two cell types flown on BioSentinel: a wild type *S. cerevisiae* strain and a *rad51* deletion mutant (*rad51*Δ) (Table **1**) [16]. The mutant is defective for recombinational repair of DNA double-strand breaks and therefore more sensitive to ionizing radiation. The position and health state of every cell is stored at each timepoint, which is what allows growth curves and growth rates to be recovered.

**FIG 1.**
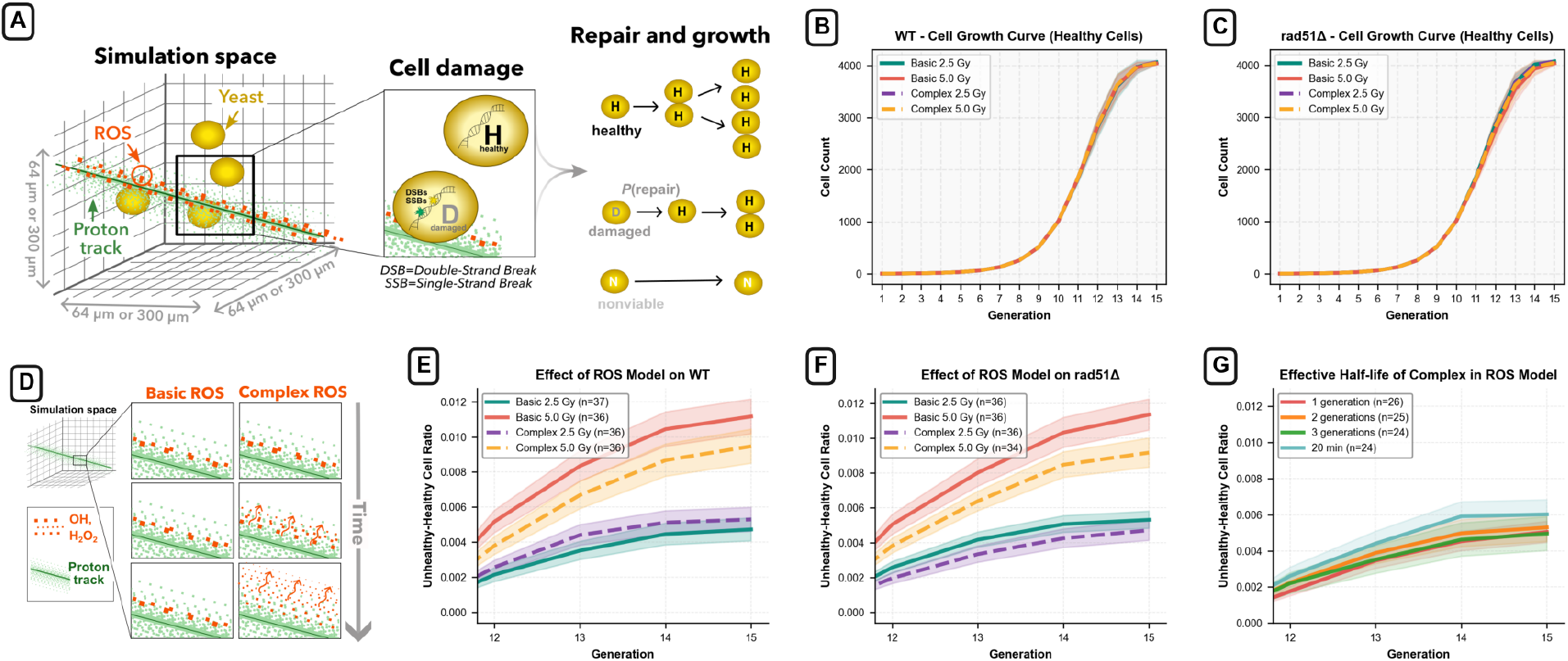
Overview of the AMMPER radiation simulation and comparison of ROS models. **(A)** Schematic of the AMMPER simulation: an ionizing proton track traverses the medium, depositing energy that generates reactive oxygen species (ROS), which damage nearby yeast cells. The diagram also depicts the cell damage, repair, and growth/replication mechanisms. **(B)** Simulated growth curves for wild-type (WT) cells under the naive ROS model at 2.5 and 5.0 Gy and the complex (diffusion-and-decay) ROS model at 2.5 and 5.0 Gy. **(C)** As in (B) but for *rad51*Δ cells. **(D)** Conceptual comparison of the naive ROS model (static, persistent molecules) and the complex ROS model (diffusion and decay). **(E)** Effect of the ROS model on WT cells, shown as the unhealthy-to-healthy cell ratio over time for the naive and complex models (same configurations as the growth curves). **(F)** As in (E) but for *rad51*Δ cells. **(G)** Effect of ROS half-life in the complex ROS model, comparing half-lives of 20, 80, 160, and 240 min (i.e., 20 min and 1, 2, and 3 yeast generations, where one generation is 80 min).

**TABLE 1.** The two *S. cerevisiae* strains modeled in AMMPER.

| Strain | Relevant genotype | Radiation-relevant phenotype |
| --- | --- | --- |
| Wild type | *RAD51* intact | Functional homologous recombination;<br>baseline sensitivity to ionizing radiation |
| *rad51Δ* | *rad51* deletion | Defective recombinational repair of double-strand<br>breaks; increased sensitivity to ionizing radiation |

### AMMPER-2

Here we describe four improvements we have recently made to AMMPER. First, we added a model of alamarBlue (aB) dye dynamics using Michaelis-Menten kinetics. Because aB is a redox-sensitive dye widely used in spaceflight experiments to study the effects of radiation on cell metabolism, this makes AMMPER’s predictions testable against metabolic as well as growth measurements. Second, we built a more accurate simulation of ROS dynamics. AMMPER-1 treats ROS molecules as static and persistent, which is fast but misses their diffusion and decay, so we added models for both and left the user free to switch between them. Third, we integrated a Graphical User Interface (GUI) to accommodate users across a broad spectrum of proficiency levels. Fourth, we made coding modifications to improve running time. We also added a gamma radiation mode, described in the Results.

### Cell metabolism and alamarBlue (aB)

AMMPER’s development was inspired by the BioSentinel and LEIA missions [35, 36, 37, 23, 24], which like several prior automated biological missions [18, 20, 19] use the metabolic dye alamarBlue (aB) as a cell health indicator [25]. Metabolically active cells generate electron carriers that irreversibly reduce the blue form of aB (resazurin) to a pink form (resorufin), which is further reduced by a reversible reaction to a colorless form (dihydroresorufin) [25, 38]. These reactions are illustrated in Figure **2**A.

**FIG 2.**
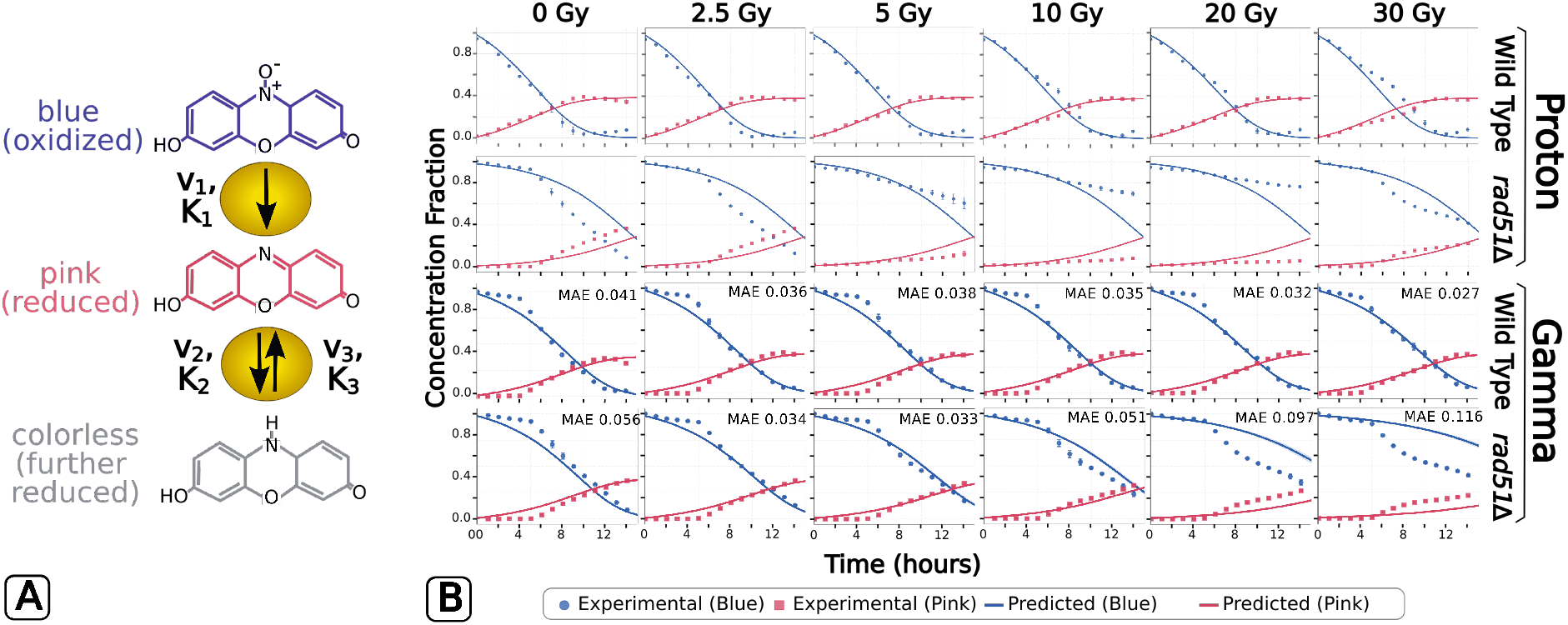
AlamarBlue (aB) reduction chemistry and Michaelis-Menten model predictions under proton and gamma irradiation. **(A)** Sequential reduction of alamarBlue: the oxidized blue form (resazurin) is irreversibly reduced to the pink form (resorufin) with maximum rate *v*_1_ and constant *K*_1_, and the pink form is reversibly reduced to the colorless form (dihydroresorufin) with *v*_2_, *K*_2_ forward and *v*_3_, *K*_3_ reverse; arrows give the direction of conversion. **(B)** Predicted and measured blue and pink aB concentration fractions at 0, 2.5, 5, 10, 20 and 30 Gy, one axes per dose. Predictions are solid lines and measurements are points; each prediction is the mean over the archived replicate simulations for that condition. The four rows are, from top to bottom, proton-irradiated wild type, proton-irradiated *rad51*Δ, gamma-irradiated wild type and gamma-irradiated *rad51*Δ. One set of kinetic constants is fitted jointly to the twelve proton conditions (rows 1 and 2), with only the inoculum offset *g*_0_ differing between strains; the two gamma rows use those constants unchanged on a separate experiment, refitting only *g*_0_ per strain, at the hit rate calibrated against the growth channel alone (Supplementary Text S2). The mean absolute error (MAE) of each gamma condition is given in its axes. In *rad51*Δ the residual is larger than in the wild type under both radiation qualities, and in both cases the measured curves flatten toward dose-dependent plateaus while the predictions continue to descend.

In these experiments cells are cultured in a microfluidic card equipped with arrays of light-emitting diodes and photodiode detectors, which measure absorbance at several wavelengths. Growth is tracked by optical density at 850 nm, and metabolism by the abundance of the blue and pink forms of aB, read at 570 nm and 630 nm. AMMPER-1 simulated the growth curve alone. For AMMPER-2 we sought to simulate aB dynamics as well. That uses the additional 570 and 630 nm data, and lets the model predict the relationship between radiation exposure and metabolic activity on a metric directly comparable with experiment. Converting those absorbances into the concentration of each aB species requires a normalization, which Supplementary Text S4 gives. It accounts for the overlap between the two species’ spectra, and for the difference between the flight LED wavelengths and the absorbance maxima at which our fitting data were taken.

## RESULTS

### AMMPER generates alamarBlue reduction curves using a naive Michaelis-Menten bulk model

We used Michaelis-Menten kinetics to construct an alamarBlue mechanics model of the reduction chemistry in Figure **2**A. It takes the growth curve produced by an AMMPER simulation as input and returns the fraction of each aB species in the medium over time. The model makes two simplifying assumptions. First, reaction rates are not distinguished inside and outside cells, so the calculation runs on population-level growth curves with no spatial information. Second, unhealthy cells consume aB at a reduced rate set by a scaling factor *k*. Treating metabolism at the population level neglects cell-to-cell heterogeneity and any spatial structure in dye reduction, and *k* represents the metabolic activity of damaged cells with a single parameter. We consider two reactions: the irreversible reduction of blue aB to pink (Equation 1) and the reversible conversion of pink to colorless (Equation 2). We estimate *K*_1_, *K*_2_, *K*_3_, *v*_1_, *v*_2_, *v*_3_ and *k* by global search against the 150 MeV proton measurements (Figure **2**B, first two rows; see Materials and Methods). Each predicted curve is the mean over the archived replicate simulations for that condition, 20 to 62 runs per dose depending on how many were archived.

In a typical aB reduction curve the blue value falls over time as yeast metabolic byproducts reduce the dye. The pink value rises before falling in turn, as it is reduced to the colorless species, which is not tracked optically. We consider only the first 15 hours of each curve, since the blue-to-pink conversion is the phase of greatest interest for the yeast experiments this model is intended to support [36, 37]. AMMPER begins each simulation from a single cell, whereas the plate-reader assay begins from a dense inoculum. We therefore introduce a single offset *g*_0_, the point on the simulated growth curve that corresponds to experimental *t* = 0. It is fit once per strain and held fixed across all six doses.

The redox chemistry of alamarBlue is a property of the dye rather than of the strain. We therefore fit one set of kinetic constants to all twelve conditions at once, six doses in each of two strains, and let only *g*_0_ differ between strains. Nine parameters then account for twelve curves, and the strain difference is carried entirely by the population state. Text S3 compares this against fitting the strains separately.

For the wild type, the predicted curves match the measurements with a mean absolute error of 0.030, ranging from 0.023 to 0.048 across the six doses (Figure **2**B, first row). The fitted offset is *g*_0_ = 8.8 generations, approximately 463 cells at *t* = 0. Refitting *g*_0_ per dose moves it by less than one generation (8.2 to 9.1), so a single shared value suffices.

For *rad51*Δ the same kinetic constants give a mean absolute error of 0.072 at *g*_0_ = 6.2 generations. The mutant population behaves like a wild type population roughly 2.6 generations behind, consistent with the slower net expansion expected of a repair-deficient strain. Its residual is concentrated at the two lowest doses (0.097 at 0 Gy and 0.091 at 2.5 Gy) rather than at the high ones, and takes a specific form. The measured curves flatten toward dose-dependent plateaus in the second half of the time course, while the predicted curves continue to descend (Figure **2**B, second row). For both strains the predictions follow the trends in the data for the blue and the pink species, with the deviations described above.

Plotted against the experimental series on matched axes (Fig. S1, Fig. S2), the predicted curves are ordered by dose in the same direction as the measured ones in both strains. The separation between doses is compressed in both. At the timepoint of greatest separation the spread across the six wild type curves is 0.057 in concentration fraction for blue and 0.022 for pink, against 0.225 and 0.103 measured. For *rad51*Δ it is 0.067 against 0.675 and 0.025 against 0.309. The model therefore recovers the direction, shape and timing of the dose response, and between a quarter and a twelfth of its magnitude.

The fitted parameters say where that magnitude is lost. The weight on damaged cells is small (*k* = 0.034), so the predicted aB signal is carried almost entirely by the healthy-cell count. That count varies by less than eight percent across the full 0–30 Gy range, because direct damage is confined to the neighborhood of the ion tracks and most cells are never intersected by one (Fig. S3). The dose dependence entering the dye chemistry is thus already small before any dye chemistry is applied. Among the rate constants, *K*_2_ and *K*_3_ are large relative to the substrate available. That places the reversible pink-to-colorless step in a first-order regime, in which these data constrain only the ratios *v*_2_/*K*_2_ and *v*_3_/*K*_3_. The constant *K*_1_, which governs the blue-to-pink step that the assay measures directly, is constrained. Text S3 reports the dose orderings, the identifiability tests and the per-condition errors.

### The aB model transfers to a gamma radiation environment

AMMPER-1 simulates only the 150 MeV proton environment. In AMMPER-2 we added a gamma radiation mode, in which the simulation volume is scanned by a plane that deposits energy at random positions rather than along a track (Fig. S4). Gamma photons deposit energy far more uniformly than charged particles do [33, 29]. This provides a test of the aB model that the proton data cannot: the kinetic constants were fit to proton conditions, and the gamma measurements are a separate experiment in which the population responds to dose much more strongly. The implementation, the calibration described below, and the full set of results are given in Supplementary Text S2.

To build the gamma model we first had to determine how many deposition events per plane a given dose represents. We calculated that number as the product of the dose and a hit rate *k*, in events per gray per plane. Because that product is all the simulator uses, choosing *k* rescales the dose axis rather than changing the model. We confirmed this directly using archived runs at a fixed nominal dose and three different hit rates (Supplementary Text S2). We calibrated *k* against the measured OD750 growth channel alone, using the ratio of the final population to the unirradiated control in both strains at all doses. That gives *k* = 1.85 events Gy^−1^ plane^−1^ (Supplementary Text S2). The alamarBlue blue and pink series that the kinetic model predicts play no part in this calibration, so the errors reported below are a consequence of it rather than a target of it.

With the dose axis calibrated, we predicted the gamma aB measurements using the proton kinetic constants unchanged, refitting only the inoculum offset *g*_0_ per strain as we did for the two proton experiments. The mean absolute error over the twelve gamma conditions is 0.049, comprising 0.035 for the wild type and 0.064 for *rad51*Δ. The same strains give 0.030 and 0.072 under protons (Figure **2**B, third and fourth rows). The gamma comparison is therefore as good as the proton one, with the dye chemistry carried across unchanged. Refitting the kinetic constants on the gamma data as well improves the mean only from 0.049 to 0.042 (Supplementary Text S2). The wild-type residual is uniform across dose (0.027–0.041). In *rad51*Δ it is concentrated at 20 and 30 Gy (0.097 and 0.116) and takes the same form as under protons. The measured curves flatten toward dose-dependent plateaus that the model does not produce.

The gamma environment also supplies the dose-dependent population response that the proton simulations do not (Supplementary Text S2). Under protons both strains retain more than 92% of the control population across 0–30 Gy. Under gamma the wild type behaves similarly (89% at 30 Gy), but *rad51*Δ falls to 33% of the control. The reason is that spatially uniform damage reaches cells that ion tracks miss entirely, and a repair-deficient strain has no intermediate damaged state in which to absorb it. The kinetic model reproduces the aB response across this much wider range of population trajectories without re-tuning. That is evidence that the compression described in the previous subsection originates in the population model rather than in the dye chemistry.

### Implementing decay and diffusion of reactive oxygen species produces little benefit over the naive model

AMMPER-2 lets the user choose a Reactive Oxygen Species (ROS) model that includes decay and diffusion (Fig. **1**D). In AMMPER-1 ROS molecules were motionless and eternal, making any location holding one a permanently toxic location, whereas in reality ROS diffuse through the medium and decompose on reacting with other molecules. We therefore use the propagator of the diffusion equation to spread the *H*_2_*O*_2_ liberated at each deposition event over the lattice points around it (Materials and Methods, Eqs. 9–11). The resulting distribution of discrete molecule counts is approximated by a multivariate normal, which is simpler to sample than the formally appropriate multivariate

Conway-Maxwell-Poisson distribution and adequate here (Fig. S5; Supplementary Text S1). *OH* remains at the deposition point in the present implementation. Parameter values follow the physical properties of ROS molecules. The decay rate depends on the chemical composition of the medium. The half-life under our experimental conditions is not known, so we took published values as a guide. These span 20 min to 5.5 hrs, with experimental values in HeLa cells at the low end, 20–30 min [39, 40, 41, 42]. Our baseline uses an intermediate half-life of 80 min, one AMMPER yeast generation.

At these baseline parameters the naive and the complex ROS models give similar damage predictions. Across multiple runs per condition, we analyzed the growth curves (Fig. **1**B,C) and the ratio of unhealthy to healthy cells over time (Fig. **1**E,F). For both cell types and all doses the complex model gave lower overall unhealthy/healthy ratios, but the two sets of curves were not statistically distinguishable (*p >* 0.05). The strain *rad51*Δ scores damage as cell death, so its ratio is the largest. There, a Cumulative Link Mixed Model (CLMM) comparison of medians found no difference between the two ROS models at any dose or timepoint. Pairwise Wilcoxon rank-sum tests at individual timepoints, run for both strains, agree: they found no significant difference at 2.5 or 5 Gy in either. Because of its greater complexity the new model also takes longer to run, with the running time scaling in the number of radiation events.

We then asked whether varying the ROS half-life would reveal an effect, scanning 20, 80, 160 and 240 min across the literature range with approximately 25 runs per condition (Fig. **1**G). A Kruskal-Wallis test across half-lives gives *p* = 0.175, so there is insufficient evidence that any half-life produced a different distribution of unhealthy-to-healthy ratios. Any effect it has is obscured by the model’s variance. The time ROS molecules persist in the medium does raise the probability of damaging nearby cells, but it matters less than where the radiation tracks fall relative to the population. The additional complexity of modeling ROS diffusion and decay therefore provides minimal advantage over the naive model under our simulation conditions. Two features of the implementation bound how large an effect either comparison could have found. First, only *H*_2_*O*_2_ is spread by the propagator, so the *OH* that produces strand breaks is not relocated relative to the naive model. Second, removal is applied once per generation, so the half-life is resolved only in units of the generation time (Materials and Methods). Extending either is the way to test whether the null result survives.

### Variance among AMMPER simulations is associated with localization of damage from ion tracks

The high variability among individual ion radiation simulations led us to investigate its source. Ion track positions are generated randomly, and radiation occurs at generation 2, when the population is four cells. We found that when the track misses the population at that point the cells are rarely damaged at all. The distribution of unhealthy-to-healthy ratios is therefore strongly zero-inflated and far from normal. Across all 380 archived simulations 115 register no radiation damage. The result is a distribution with a point mass at zero, a short right tail and no interior mode. Even at the highest dose the mean ratio of damaged to healthy cells stays below 0.07 in both strains (Fig. S3A, Fig. S3B). This shape is not arbitrary. It follows from the generative structure of the simulation, in which a track either intersects the four-cell population or misses it. A parametric zero-inflated count description would be expected to fit it. We do not fit one here, and use distribution-free tests instead (Materials and Methods). Most of those zeros are the unirradiated controls, which record no damage by construction (92 runs). The informative part is that 23 of the 288 irradiated runs also record none. All 23 fall at 2.5 or 5 Gy, where the track count is lowest and a miss is correspondingly likely. This happens more often in the complex ROS model. There, ROS can only damage cells in early generations, before diffusion and decay reduce the local concentration, whereas in the naive model persistent molecules can still damage cells that encounter them late. GCR particles traverse a yeast nucleus in deep space only rarely, approximately once every 60 days for protons and *α* particles behind 1 g cm^−2^ of aluminum at solar minimum [43], with traversals by heavier nuclei rarer still. A zero-inflated damage distribution may therefore be accurate. Further work is needed on how best to structure AMMPER to reproduce the true distribution.

The corollary is a limitation of the comparison between radiation modes. Because ion tracks miss most cells, the proton simulations deliver their nominal dose to a small and randomly selected part of the population, whereas the gamma mode deposits energy uniformly across the volume and therefore reaches every cell. The two modes are matched in nominal dose but not in how many cells that dose reaches, which is why *rad51*Δ loses two thirds of its population under gamma at 30 Gy and less than a tenth under protons. That ordering should not be read as a statement about the relative biological effectiveness of the two radiation types, which for low-LET protons is close to unity relative to photons [44]. It is a consequence of how the two modes were set up: the gamma hit rate was calibrated against measured survival, whereas the proton track count is a fixed table. Only one of the two modes is therefore tied to a measured population response.

### Graphical user interface enhances ease of use

To make deep space radiation simulations accessible to a broader audience, we built a graphical user interface (GUI) that replaces the sequential command-line workflow of AMMPER-1 with a single interactive dashboard (Fig. **3**). The GUI is written in Python and built on PyQt5 (version 5.15.9). Simulation output is rendered in the window with Matplotlib (version 3.7.2). The animated slideshows are assembled with MoviePy (version 1.0.3). The full dependency list is given in the AMMPER repository. Where AMMPER-1 required users to answer command-line prompts one at a time and then locate output files by hand (Fig. **3**A), the GUI removes the need to read or modify Python code. A start screen offers either interface. The dashboard collects the simulation parameters (cell type, radiation dose, simulation type, ROS model and output directory), runs AMMPER, and renders its outputs in the same window (Fig. **3**B). Beyond the outputs of the base package it also produces dynamic visualizations as kinetic icons and slideshows, so a run can be followed visually over time. The command-line interface remains preferable for many replicate iterations. The GUI serves AMMPER’s mission of providing modeling tools to researchers in microbial radiobiology for whom modeling might otherwise be out of reach, and an interactive resource for space biology education [45, 46, 47, 48].

**FIG 3.**
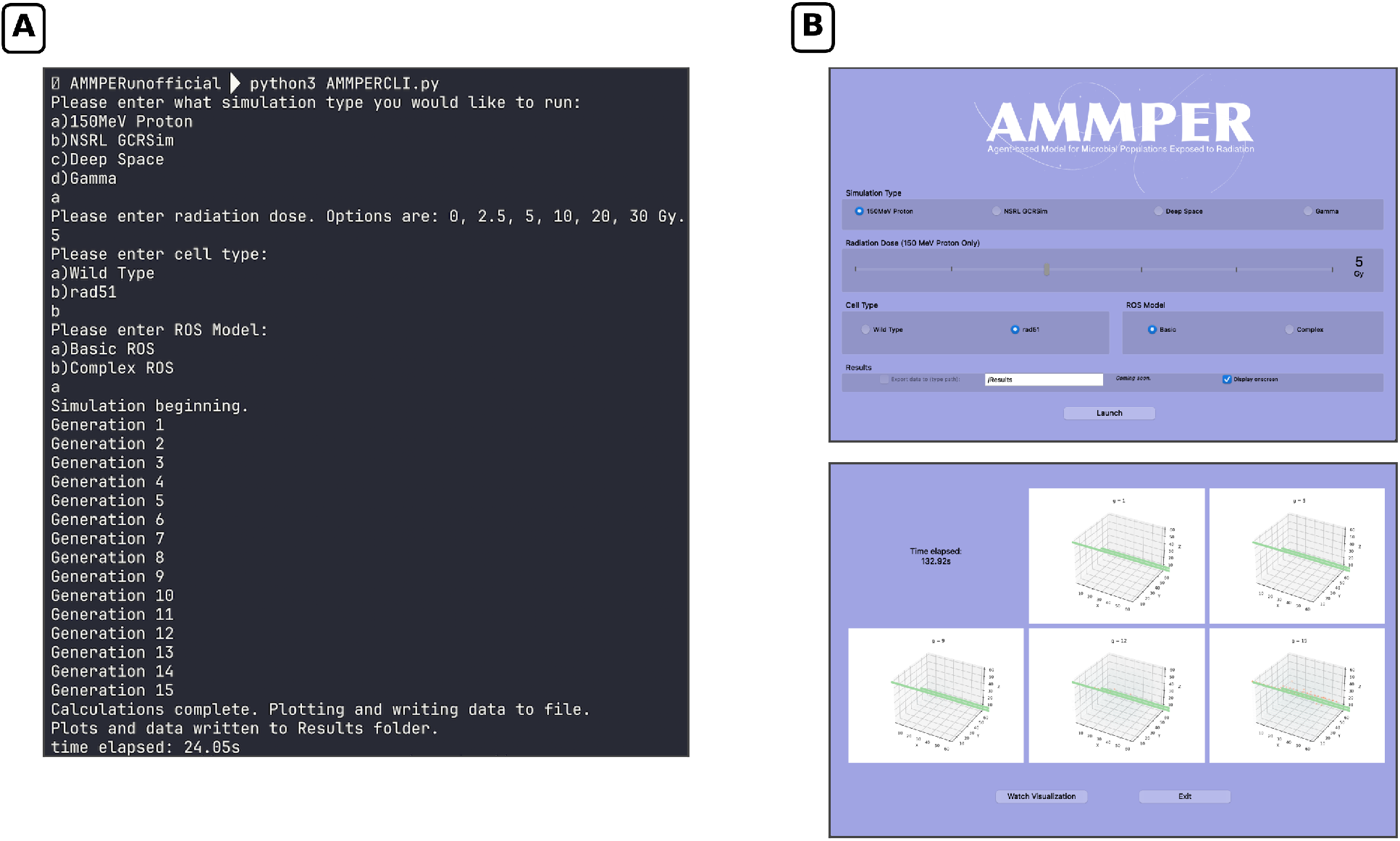
Workflow comparison between the original AMMPER (AMMPER 1) interface and the redesigned AMMPER 2 GUI. **(A)** AMMPER 1 workflow, in which parameter selection, simulation execution, and results retrieval occurred as separate, sequential steps: users answered CLI prompts one at a time, then manually located output files after the run completed, with no visual feedback during execution. **(B)** AMMPER 2 workflow, in which all simulation parameters (simulation type, radiation dose, cell type, ROS model, output path) are set from a single configuration screen, launched with one click, and monitored in real time via on-screen rendering of simulation state at successive generations.

### AMMPER-2 requires reduced computational resources due to optimization of visualization process and ROS damaging mechanics

AMMPER is designed to run on a standard desktop computer, so in AMMPER-2 we sought greater computational efficiency than AMMPER-1. Two changes account for most of it. Visualizations now plot all cells at once per generation (Algorithm 1C) rather than cell by cell (Algorithm 1A). That reduces the memory overhead that was the main bottleneck, giving *O*(*G*) plotting calls instead of *O*(*G · N*) for *G* generations and *N* cells. ROS damage is now evaluated per cell over the events in its neighborhood (Algorithm 1D) rather than per event over all cells (Algorithm 1B). That reduces *O*(*R* · *N*) for *R* events to approximately *O*(*N* · *k̅*), for *k̅* the mean number of events near a cell. The original approach became impractical once the diffusion model spread ROS more widely through the volume.

#### Algorithm 1

Pseudocode illustrating the two changes from AMMPER-1 that account for most of the performance improvement in AMMPER-2. Parts A and B give the AMMPER-1 implementations. Parts C and D give the AMMPER-2 replacements

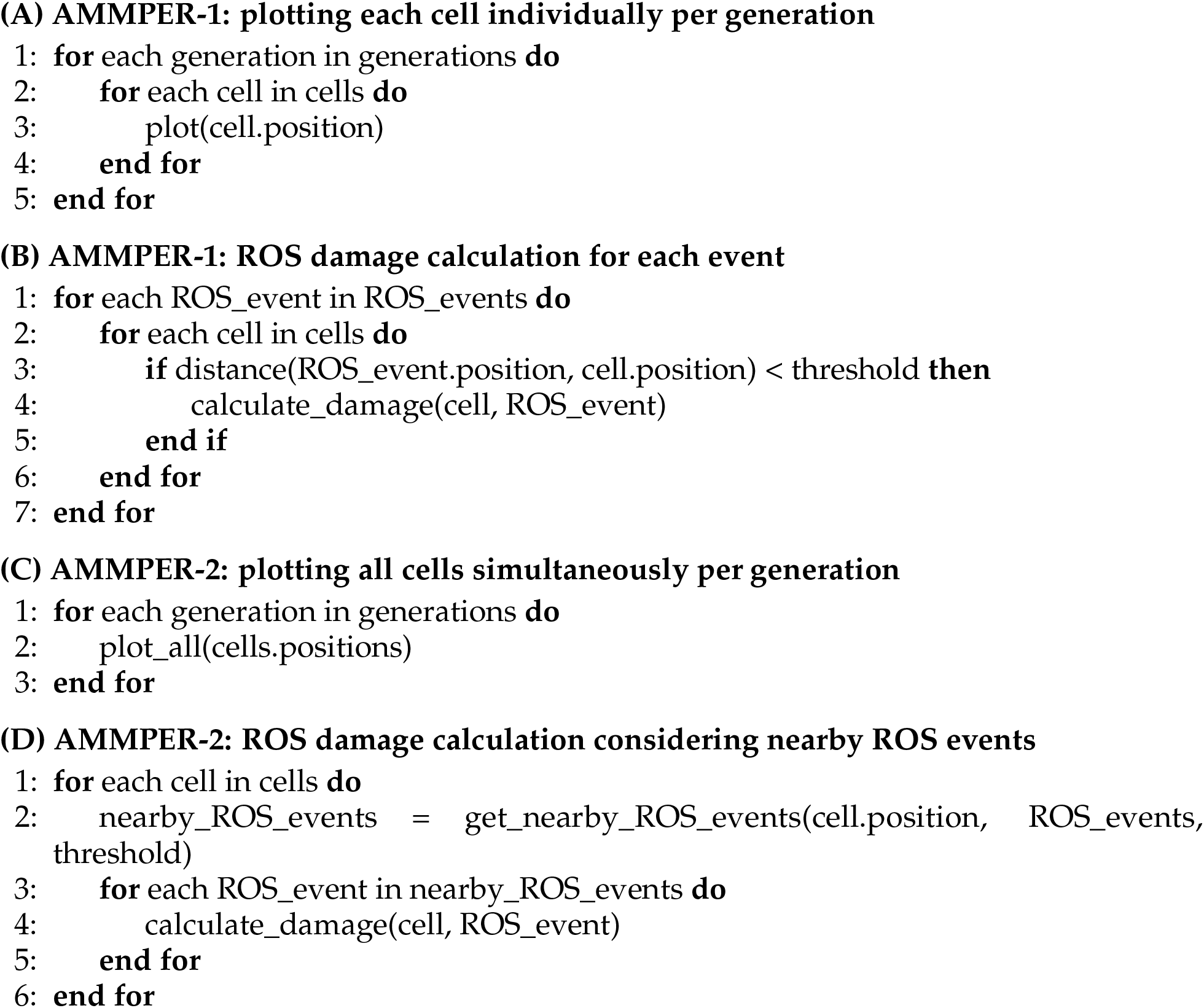

We measured the second change directly, running both implementations on identical inputs and confirming that they damage the same cells (Fig. S6). The spatially filtered implementation is faster above approximately 1,000 ROS lattice points, by a factor that grows with problem size and reaches 8.0× at 8,000 points. Below that it is slightly slower, because the fixed cost of a vectorized filter is not amortized over few points. The optimization therefore matters precisely in the regime the diffusion ROS model creates, since that model emits 82 lattice points per deposition event where the naive model emits one.

## DISCUSSION

### Strengths of the AMMPER model

AMMPER’s first-order strength is that it reproduces population-level measurements of the kind generated by past and future flight missions. Here those measurements are growth and alamarBlue reduction, in both BioSentinel strains and under two radiation qualities. It reproduces them from an explicit model of damage to individual cells, and so connects two scales that flight experiments cannot themselves bridge. A second strength is low computational cost and the accessibility that follows from it. With default settings AMMPER runs on a standard laptop, and the GUI makes it usable without coding experience, which matters for a tool intended for researchers and educators as much as for modelers. Its modularity is a related strength, in that individual components can be modified independently. The lifetime-diffusion ROS model presented here is one such component, and with appropriate parameters the naive model reproduces the damage it predicts. Adjusting the initial parameters (number of initial cells, simulation length and volume, timing of radiation) also lets AMMPER address experiments beyond BioSentinel and LEIA.

The aB model itself is simple, and this is a further strength. Simulating radiation damage at a metabolic level would conventionally require chemical agents, metabolic networks and a mechanism coupling radiation damage to them. The aB model instead reaches a mean absolute error of 0.030 in the wild type with nine parameters, and it reproduces the trends in the experimental data (Figure **2**B and Fig. S1, Fig. S2). It also extends to a second strain and to a second radiation quality without re-tuning the dye chemistry.

### Limitations and future directions

The clearest limitation is the AMMPER cell health mechanics, which distinguish only healthy, damaged and dead states. This is what limits the model’s dynamic range. The predicted curves are ordered by dose in the correct direction throughout and, for *rad51*Δ, perfectly monotonically. What is compressed is the magnitude of the separation, roughly fourfold to fivefold in the wild type and tenfold or more in the mutant (Fig. S1, Fig. S2). Because the aB model takes the population trajectory as its only input it cannot spread more widely than that trajectory allows, and the simulated healthy-cell count varies by under eight percent across the full 0–30 Gy range. A damaged *rad51*Δ cell is moreover assigned directly to the nonviable state, so that strain produces no intermediate-state cells at any dose, and the fitted weight on damaged cells is small in any case. Expanding the damaged state to represent modified metabolism — a graded health state with an explicit per-cell metabolic rate — would let dose modulate the aB signal through the metabolic activity of surviving-but-impaired cells rather than through the healthy-cell count alone. The residual misfit in *rad51*Δ has a form consistent with this: the measured curves flatten toward dose-dependent plateaus that a model without a graded, dose-sensitive metabolic state cannot produce.

The gamma mode carries its own limitations. It has been calibrated against a single published survival dataset [49] and one alamarBlue experiment per strain, so its hit rate should be read as fit to those measurements rather than derived from radiation transport. The calibrated rate also differs between strains when they are fit separately (6.84 against 1.85 events Gy^−1^ plane^−1^), which a geometric property of the radiation should not. We report the shared value and note the discrepancy in Supplementary Text S2. The gamma mode is also where the binary health state is most visible. The residual misfit concentrates in *rad51*Δ at the two highest doses, exactly where uniformly distributed damage should leave many surviving-but-impaired cells that the implementation cannot represent. Attenuation and shielding are neglected, which is defensible at the 64^3^ *µ*m^3^ scale of the simulation volume but would not be at a larger one.

A graded health state and additional radiation environments are the directions in which we expect a more complete account of cellular dynamics under deep space stress to be found.

## MATERIALS AND METHODS

### Microbiological methods and radiation exposure experiments

Empirical data on yeast growth and metabolism under proton irradiation used in this report have been previously reported in full [37, 32]. The gamma irradiation series is reported here for the first time; it was generated by the same group using the protocol described below. Briefly, the yeast strains used were diploid prototrophic derivative of the W303 background (*MAT*a/*MATα ADE2 CAN1 HIS3 LEU2 TRP1 URA3*). Standard genetic cross and tetrad dissection procedures were used to generate the *rad51*Δ strain from the wild type. In preparation for irradiation, samples were grown in liquid Yeast extract-Peptone-Dextrose (YPD) broth at room temperature (*∼*23^◦^C) for 7 days, then washed and resuspended in sterile water with 10% trehalose, deposited into 96-well stripwell microplates (9102; Costar®) at a density of 10^5^ viable cells per well, and air-dried for at least 7 days. Cells were rehydrated with 100 *µ*L of Synthetic Complete growth medium containing 1*×* aB (DAL1100; Invitrogen) immediately prior to radiation exposure. Proton exposure was conducted at Loma Linda Proton Treatment and Research Center using a 20-cm diameter circular unmodified beam; gamma exposure was conducted at NASA Ames Research Center inside a ^137^Cs source chamber. After exposure, cell suspensions were transferred to clear 96-well nontreated polystyrene flat-bottom microplates (351172; Falcon®) and grown at room temperature in Molecular Devices SpectraMax or VersaMax plate readers with absorbance measurements taken at 570 nm, 600 or 630 nm, and 750 nm once per hour for at least 48 hours. Absorbance measurements at these wavelengths were used to calculate growth via optical density at 750 nm, and alamarBlue reduction state, as described in Supplementary Text S4.

### AlamarBlue metabolism Michaelis-Menten irreversible kinetics model

Yeast metabolism at the genome level is complex, with thousands of reactions, co-factors, metabolites, genes, and other components [50, 51, 52, 53, 54]. It is not clear how ionizing radiation affects each individual component of the network; deleting individual genes or components would likely miss the physically disruptive effect of energy deposition by an ion track. We therefore focused AMMPER on modeling alamarBlue (aB) metabolism at each individual cell as being dependent on the overall cell health state of the whole cell. In order to make the aB uptake rate of each cell concentration-dependent, Michaelis-Menten kinetics [55, 56, 57] are used:

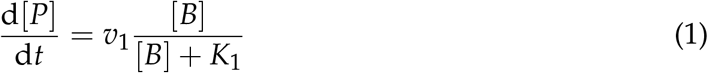

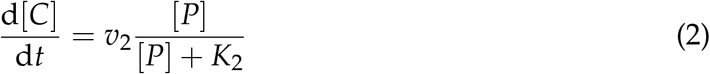

Where [*P*] denotes the pink (aB reduced), [*B*] the blue (aB oxidized), and [*C*] the clear concentrations. Here, *v*_1_, *v*_2_ are the maximum rates, and *K*_1_, *K*_2_ are the equilibrium constants. Then, the AMMPER growth curves are used to update the total intake rates per generation:

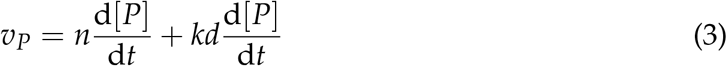

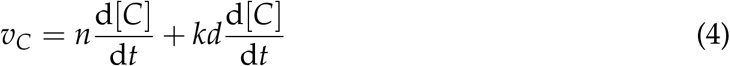

Where *n* and *d* correspond to the total number of healthy and damaged cells for a given generation. The intake rates and concentrations are calculated recursively and are generation dependent. Here, *k* serves the same purpose as in the preliminary models described above. Each concentration is normalized by dividing by the total concentration. Because the kinetic constants are properties of the dye rather than of the radiation exposure, one set is fitted jointly to all twelve conditions (six doses in each of two strains). Only the inoculum offset *g*_0_ is allowed to differ between strains. The search procedure is described under “Parameter fitting for the kinetic model” below. We implemented this model in Python with supporting packages such as numpy, scikit-learn, and scipy.

### AlamarBlue metabolism Michaelis-Menten reversible kinetics model

The process from pink to clear is reversible, so Equation 2 requires a small modification to represent it. We use the notation found in the literature [58]:

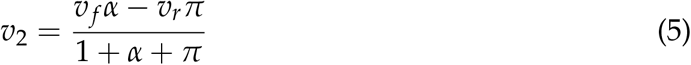

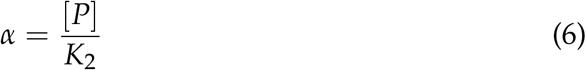

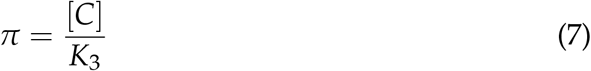

This introduces a new pair of parameters that we rename for convenience:

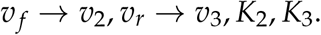

### Error metric

To assess how well the fitted model recapitulated the experimental data, the experimental and predicted curves were compared using the mean absolute error. The error metric is the mean of the absolute deviations between predicted and experimental values, averaged over both aB species and all measured timepoints:

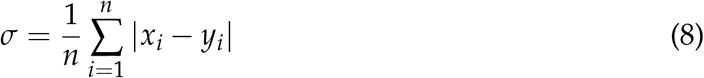

This is the objective minimized during fitting and the metric reported throughout the Results and in Supplementary Text S3. The colorless species is not measured optically and therefore does not enter the error independently. The concentrations are normalized so that [*B*] + [*P*] + [*C*] = 1. The deviation in [*C*] is therefore fixed by those in [*B*] and [*P*], and scoring it separately would double-count the same discrepancy.

### Parameter fitting for the kinetic model

The rate constants reported in the Results were estimated by fitting the kinetic model to all twelve experimental conditions at once (six radiation doses in each of two strains). Because the redox chemistry of alamarBlue is a property of the dye rather than of the genotype, the six rate constants and the weight *k* on damaged cells are shared across all twelve conditions. Only the inoculum offset *g*_0_ differs between strains, which gives nine parameters for twelve measured curves. We used differential evolution (scipy.optimize.differential_evolution, seeded for reproducibility) as a global search, and refined each offset by exhaustive scan with the kinetics held fixed. The objective is the mean absolute error of Equation 8, computed over both species and all measured timepoints. The choice of optimizer is not itself under test; the same objective could be minimized by other means. The bounds and the random seed are recorded with the fitted values, and the search is described in full in Supplementary Text S3.

Predictions and measurements are placed on a common scale before the error is computed, so that the reported values are interpretable as fit quality rather than as a units mismatch. The experimental series are normalized against the total dye present at *t* = 0, so the measured blue and pink fractions together with the unmeasured colorless remainder sum to unity. The kinetic model conserves dye by construction, so its three species sum to unity at every timepoint. Text S3 gives the checks on this: that the implied colorless fraction and the measured blue-plus-pink sum behave as the reaction chemistry requires, and that no part of the reported discrepancy is attributable to the two series being scaled differently.

The resulting values are *v*_1_, *v*_2_, *v*_3_ = 3.22, 18.6, 16.0 and *K*_1_, *K*_2_, *K*_3_ = 1.39 *×* 10^4^, 218, 303, with *k* = 0.034, *g*_0_ = 8.81 generations for the wild type and *g*_0_ = 6.21 for *rad51*Δ. We note that *K*_2_ and *K*_3_ are both large relative to the amount of the corresponding substrate present. That places the reversible pink-to-colorless pair in an effectively first-order regime, in which only the ratios *v*_2_/*K*_2_ and *v*_3_/*K*_3_ are constrained by these data. Scaling either pair together by a factor of 100 leaves the mean absolute error unchanged to three decimal places, whereas the same scaling applied to (*v*_1_, *K*_1_) nearly doubles it. *K*_2_ and *K*_3_ should therefore be regarded as not individually identifiable from the present measurements, and we do not interpret their absolute magnitudes. *K*_1_ is constrained. The per-condition errors, the identifiability checks, and the treatment of mass conservation in the integrator are given in Text S3.

### Diffusion and decay ROS molecule modeling

AMMPER-1 modeled ROS molecules as maintaining static locations in space and existing eternally. This simplification reduced the model’s complexity and decreased the computational time for each simulation, but it is not physically accurate: real ROS molecules move away from where they were created and are consumed over time. AMMPER-2 therefore replaces this treatment with two explicit processes, diffusion and decay, described in turn below. Both act on the individual ROS molecules already generated by the RITRACKS energy deposition events, and both matter because the damage a cell sustains depends on how close ROS molecules come to it and for how long.

#### Diffusion

The physical starting point is the diffusion equation,

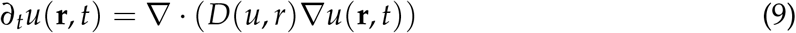

where the vector **r** is the location in three-dimensional space, *D* is the diffusion coefficient, and *u* is the density of the diffusing material. Each energy deposition event is effectively a point source, so the relevant solution of Equation 9 is its Green’s function, or propagator [59, 60]:

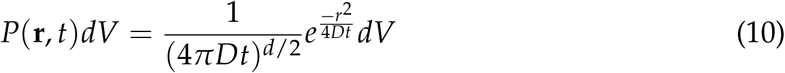

Equation 10 gives the probability of finding a molecule a distance *r* from its point of origin after a time *t*, and it is a Gaussian whose width grows as 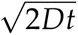. This is what makes the implementation straightforward: rather than integrating Equation 9 on the AMMPER lattice, we can distribute the ROS liberated at each deposition event over the lattice points around it in proportion to the corresponding multivariate normal density,

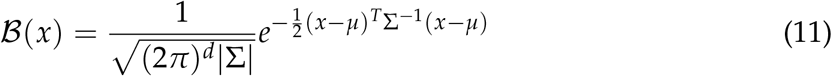

where the mean *µ* is the location of the deposition event and the covariance matrix Σ sets the spread. We implemented Equation 11 with the SciPy Python package [61], evaluating it at each deposition event on a 9 *×* 9 neighborhood of lattice points and distributing the *H*_2_*O*_2_ yield over that neighborhood in proportion to the density. Three properties of the present implementation should be stated plainly, since they bound what the diffusion model can do. First, the covariance is fixed at Σ = [[4, *−*1], [*−*1, 4]] in lattice units rather than being set from a species diffusion coefficient and an elapsed time. The spread therefore corresponds to a single characteristic diffusion length: a standard deviation of two lattice units, or approximately 2 *µ*m at AMMPER’s 1 *µ*m lattice spacing, chosen to be of the order reported for *H*_2_*O*_2_. It does not broaden between generations. Second, the neighborhood is two-dimensional, spread in *y* and *z* with the deposition coordinate in *x* unchanged. Third, only *H*_2_*O*_2_ is spread, with the *OH* yield left at the deposition point, so the two species are not given independent propagators. The consequence is that the diffusion model changes where *H*_2_*O*_2_ is found and, through the removal process described next, for how long any molecule persists, but it does not relocate the *OH* that drives strand-break damage in either strain. This is one reason the comparison against the naive model reported in the Results finds so little difference, and giving *OH* its own propagator is the most direct extension of the present implementation. Strictly speaking, because AMMPER tracks discrete molecules on a Cartesian lattice, the per-voxel molecule counts produced by superimposed point sources follow a multivariate Conway-Maxwell-Poisson distribution rather than a Gaussian. That distribution is difficult to implement and to sample from, and Equation 11 is the continuous approximation to it that reproduces the same diffusive broadening (Supplementary Text S1).

#### Decay

ROS molecules are removed from the simulation according to an exponential decay law. At each cell replication generation after the exposure generation, the code deletes a randomly selected subset of the existing ROS molecules. The surviving fraction 2^−Δt/t^_1/2_ is set by the assumed half-life *t*_1/2_ and the generation time Δ*t*. The removal is therefore per generation rather than continuous, so the half-life enters the simulation only through that fraction, and the resolution of the scan below is limited by the generation time. The baseline is a half-life of one generation, a surviving fraction of 0.5. Because ROS decomposition rates are context-specific and depend on the chemistry of the cell medium, no single half-life is correct for our conditions. We therefore conducted a literature review to establish the physically plausible range to test in AMMPER [62, 63, 64, 65, 66], and treated the half-life as a parameter to be scanned over that range (Results).

### Statistical analysis

Two comparisons in this work are inferential, and both concern the ROS model: whether the naive and the diffusion-decay implementations give different damage, and whether the ROS half-life matters. Both were run in R (version 4.3.1) on the per-run ratio of unhealthy to healthy cells, using ordinal [67], RVAideMemoire, emmeans and rstatix on top of tidyverse.

Both comparisons use distribution-free tests by necessity rather than by preference. The damage ratio is zero-inflated. No damage is recorded in either of the unirradiated controls. Among irradiated runs the ion track is placed at random when the population is four cells, so runs in which it misses the population record no damage either (Fig. S3). Shapiro-Wilk tests within dose and timepoint groups confirmed non-normality. We therefore did not use the repeated-measures ANOVA that the design would otherwise suggest.

For the naive versus diffusion-decay comparison we binned the damage ratio into twelve ordered levels of width 0.002. We then fitted a cumulative link mixed model (clmm) with the ROS-model-by-dose combination and timepoint as interacting fixed effects and run identity as a random effect. The fixed effects were tested by type-II analysis of deviance (Anova.clmm), followed by Tukey-adjusted pairwise comparisons of the estimated marginal means. Binning to an ordinal response, rather than modeling the ratio directly, is what accommodates the point mass at zero. The model was fitted on *rad51*Δ, where damage is scored as death and the ratio spans the widest range; the same contrasts were checked in both strains at individual timepoints with unpaired Wilcoxon rank-sum tests. For the half-life scan (20, 80, 160 and 240 min) we used a Kruskal-Wallis test across half-lives, with Holm-corrected pairwise Wilcoxon rank-sum tests to follow if it reached significance; it did not, so no pairwise tests are reported. The complete outputs of both analyses are in the AMMPER repository: the model fit, its type-II ANOVA table, the marginal-mean contrasts, the rank-sum and Kruskal-Wallis results, and the Shapiro-Wilk tests used to select them. The repository also holds the R and Python scripts that produce them and every figure reported here, so no value in this work is one that cannot be recomputed.

Figures were produced in Python with Matplotlib [68] through the PubliPlots library [69]. PubliPlots fixes the size of the plot axes rather than of the canvas, so panels of the same nominal shape are physically the same size across figures. The simulation outputs and the analysis code behind every figure and statistic reported here are in the AMMPER repository (see Data Availability).

### Use of artificial intelligence

AMMPER is an agent-based simulation built on explicit kinetic and stochastic rules; no machine-learning or generative model forms any part of its implementation, and none was used to produce or analyze the results reported here. No AI-assisted tool was used for study design, analysis, or primary manuscript drafting. Generative AI tools were used only for proofreading and typographical and grammatical correction of author-written text. No AI tool is listed as an author, and the authors reviewed and verified all text and take full responsibility for the integrity, accuracy, and originality of all content.

## ACKNOWLEDGMENTS

D.P. contributed to the majority of the model development, writing, and figures. P.S. made significant contributions to overall enhancements of the AMMPER code, compiled the figures into panels, and provided revisions to the manuscript. M.M. contributed to the development of the graphical user interface (GUI) of AMMPER. D.M. contributed to the AMMPER code, including diagnosis and resolution of issues, and improvements to documentation and to the GUI. L.C.L., D.M.G., and S.R.S.M. as part of the BioSentinel team, contributed the empirical data from the experiments described in this study, as well as essential space biology expertise. J.L. conceptualized the project, provided oversight and mentorship, and edited the writing and figures.

## DATA AVAILABILITY STATEMENT

All data, source code, and models generated or analyzed during this study are publicly available. The complete AMMPER-2 source code is openly available in the AMMPER repository at https://github.com/nasa/AMMPER. It includes the alamarBlue kinetics model, the reactive oxygen species diffusion-and-decay model, and the graphical user interface, together with the simulation outputs, experimental datasets, and the analysis scripts used to generate the figures and statistics reported here. No restrictions apply to the availability of these materials.

## CONFLICTS OF INTEREST

The authors declare that they have no competing interests, financial or otherwise, that could have influenced the work reported in this manuscript. The funders had no role in the study design; in the collection, analysis, or interpretation of data; in the writing of the manuscript; or in the decision to submit the work for publication.

### Supplemental material

The supplemental file (compiled separately) contains four supplementary text sections. Supplementary Text S1 gives the multivariate distribution used for ROS diffusion. Supplementary Text S2 covers the gamma radiation mode, its hit-rate calibration, and the alamarBlue comparison under gamma exposure. Supplementary Text S3 describes fitting the alamarBlue kinetics model, and Supplementary Text S4 the conversion of absorbance measurements to aB concentrations. The file also contains Supplementary Figures S1–S9. Each item is cited from the text, and each supplementary text section names the main-text section that refers to it.

