## Supplemental File 1 for "AMMPER-2: A spatially explicit agent-based model of microbial radiobiology with redox dye simulation"

### Supplementary Material

#### Supplementary text

##### S1. Multivariate distribution used for ROS diffusion

*Corresponds to Materials and Methods, "Diffusion and decay ROS molecule modeling" in the main text, where this section is cited.*

When an ionizing radiation path is created in AMMPER using RITRACKS, a radiation track is generated in three-dimensional space, and the energy depositions that create reactive oxygen species (ROS) are single point events along that track. Each radiation event therefore acts as a point source of diffusion, and its spread is described by the propagator (Green's function) of the diffusion equation [1, 2]:

$$P(\mathbf{r}, t)\,\mathrm{d}V = \frac{1}{(4\pi D t)^{d/2}}\, e^{-r^2/(4Dt)}\,\mathrm{d}V, \tag{S1}$$

where $\mathbf{r}$ is the position, $r = |\mathbf{r}|$, $D$ is the diffusion coefficient, and $d$ is the spatial dimension. Equation S1 depends only on radial distance and time; its broadening over time is illustrated in Fig. S5.

Because AMMPER is defined on a Cartesian three-dimensional lattice and ROS are tracked as discrete molecules, the formally appropriate description of the per-voxel counts generated by superimposed point sources is a multivariate Conway-Maxwell-Poisson distribution [3, 4, 5, 6, 7]:

$$\mathcal{P}(x : \lambda) = e^{-\sum_{i=0}^{d} \lambda_i} \prod_{i=1}^{d} \frac{\lambda_i^{x_i}}{x_i!} \sum_{j=0}^{\min(x_i)} \left( \prod_{i=1}^{d} \binom{x_i}{j} \right) j! \left( \frac{\lambda_0}{\prod_{i=1}^{d} \lambda_i} \right)^j . \tag{S2}$$

In practice, Equation S2 is difficult to implement and to sample from, and no well-supported implementation was available in our software stack. We therefore approximated the spread of ROS around each deposition event with a multivariate normal (Gaussian) distribution, for which mature support exists in SciPy [8]:

$$\mathcal{B}(x) = \frac{1}{\sqrt{(2\pi)^d |\Sigma|}} e^{-\frac{1}{2}(x-\mu)^\intercal \Sigma^{-1} (x-\mu)}, \tag{S3}$$

where $\mu$ is the mean and $\Sigma$ the covariance matrix. This is the distribution reported in the main text. The Gaussian is a continuous approximation to the discrete counts described by Equation S2. It reproduces the same diffusive broadening (Fig. S5) at a small fraction of the implementation and sampling cost. In the present implementation $\Sigma$ is fixed rather than set from a diffusion coefficient and an elapsed time. The spread is taken over two of the three lattice axes, and it is applied to $H_2O_2$ only, with $\cdot OH$ left at the deposition point. The main text states what each of these bounds is. Molecules are removed once per generation according to an exponential decay law whose half-life is drawn from the experimentally reported range [9, 10, 11, 12, 13, 14, 15, 16, 17].

##### S2. Gamma radiation mode

*Corresponds to Results, "The aB model transfers to a gamma radiation environment" in the main text, where this section is cited.*

Gamma rays are an additional type of ionizing radiation found in deep space, composed of high-energy photons rather than particles. They interact with matter primarily through the Compton effect, and like galactic cosmic radiation (GCR) they can cause cell death and mutations [18, 19]. Their interaction with biological material differs from that of GCR in the localization of damage. The charged particles composing GCR produce fewer tracks with higher localized energy deposition, whereas the photons of gamma radiation deposit energy more uniformly throughout the medium [20]. This contrast motivated us to explore whether AMMPER's damage mechanics could be extended to a non-localized radiation environment.

AMMPER-1 does not simulate gamma radiation. In AMMPER-2 we implemented a gamma radiation mode as follows. Because energy deposition from gamma radiation is non-localized, a uniform probability distribution is used to determine the chance of an interaction, rather than the track-following approach used for protons. We scan the three-dimensional volume with a plane perpendicular to the direction of radiation, and each plane is assigned a probability of generating $m$ radiation deposition events. The number of events is set by a hit rate $k$, in deposition events per gray per plane, and is calibrated to reproduce survival rates from published gamma radiation experimental data [21]. The plane traverses the whole cell medium equally. The resulting difference in damage topology is visible in the AMMPER visualizations: ion tracks produce localized ROS generation along the track (Fig. S4b), whereas gamma events are distributed randomly and cell damage is non-localized (Fig. S4a).

When gamma radiation passes through a medium it is attenuated by interaction with matter and by solid-angle effects, so that cells closer to the source receive more damage and partially shield those behind them [22, 23]. AMMPER models cell populations in a $64^3$ $\mu m^3$ cube. Even at low doses the penetration length is far larger than this, on the order of $\approx$6 cm. Shielding effects are therefore neglected in the current implementation and could be added in future work [24, 25, 26, 27, 28].

**Calibrating the hit rate.** The number of deposition events per plane is $m = \lfloor Dk \rceil$ for a dose $D$ in gray. The hit rate $k$ therefore fixes the correspondence between the nominal dose label of a simulation and the physical exposure it represents. The value $k = 100$ carried by the simulator was never calibrated against a measurement, and it is 54 times larger than the value we obtain below. Its consequence is visible at 20 Gy: it puts the simulated population at 1.3% of the unirradiated control, where the measurement is at 76%. The population trajectory reaching the kinetic model has therefore gone extinct before the measured dose range begins, and no dye chemistry downstream of it could reproduce the measurements. Fixing $k$ is a precondition for the comparison, not a refinement of it. We calibrate $k$ against the OD750 growth channel. For each strain we take the ratio of the final population to the unirradiated control, in the measurements and in the simulations. We then choose the single $k$ that minimizes the root-mean-square residual in log survival across both strains and all doses. This gives $k = 1.85$ events $Gy^{-1}$ $plane^{-1}$. Fitting each strain separately gives 6.84 for the wild type and 1.85 for *rad51*$\Delta$, so the shared value is set by the mutant. The mutant is the strain whose survival actually varies over the measured range. We report the shared value and note the spread rather than fitting the two independently, since the hit rate is a property of the radiation geometry and not of the genotype.

Two things make this a calibration rather than a change to the model. First, the number of events depends on $D$ and $k$ only through their product, so choosing $k$ is exactly a rescaling of the dose axis and requires no modification to the simulator. Three archived runs at the same nominal 2.5 Gy with $k = 50$, 100 and 1000 confirm this degeneracy directly. They fall on the survival curve traced by the $k = 100$ dose series, at the events-per-plane value each corresponds to (observed survival 0.795, 0.657 and 0.006 against 0.785, 0.683 and 0.005 predicted from the dose series). Second, $k$

is fit to the growth channel alone, so the aB errors are a consequence of the calibration and not a target of it.

**Transferring the dye chemistry.** The constants transferred to the gamma prediction are the same $v_1, v_2, v_3, K_1, K_2, K_3$ and $k$ reported in the main text, with only the inoculum offset $g_0$ refit per strain. Three details of that run are specific to the gamma experiment. First, the gamma experiments run over a slightly longer window than the proton ones, so the generation time is 205.62 min here against 198 min for the protons. This is set by the experiment duration and the number of simulated generations, not fitted. Second, twelve replicate simulations are used for each of the ten irradiated conditions rather than three. At the calibrated rate the number of events per plane is small (5 at 2.5 Gy, 56 at 30 Gy), so which cells are hit is correspondingly stochastic. Third, the two 0 Gy conditions reuse the unirradiated proton archives (62 runs for the wild type, 30 for *rad51*Δ), since the simulator applies no radiation at zero dose and the two radiation modes are then identical.

The resulting per-condition and per-strain errors, and the comparison against the proton errors, are reported in the main text (Results, "The aB model transfers to a gamma radiation environment"; main-text Fig. 2B, third and fourth rows). They are not repeated here. The fitted offsets, which are not given there, are $g_0 = 7.91$ generations for the wild type and 7.62 for *rad51*Δ, corresponding to approximately 244 and 207 cells at $t = 0$.

Two comparisons place those errors. Run at the uncalibrated hit rate $k = 100$, with everything else held the same, the mean absolute error is 0.172. That is 3.5 times larger overall, 4.9-fold in the wild type and 2.7-fold in *rad51*Δ, and larger at every irradiated dose in both strains. The only exception is the unirradiated mutant control, where no radiation is applied and the two configurations differ solely through the fitted inoculum offset. Refitting the kinetic constants on the gamma data instead of transferring them improves the mean only from 0.049 to 0.042. Together these fix what the gamma result does and does not rest on. It rests on the hit rate, and it does not rest on refitting the dye chemistry. That is why we report the transferred configuration as a test of the fitted chemistry on data that did not inform it, rather than as a second fitting exercise.

**Where the residual sits.** The main text reports where the error is concentrated. It is uniform across dose in the wild type, and confined to the two highest doses in *rad51*Δ, where the measured curves flatten toward dose-dependent plateaus that the predictions do not reproduce. That is the dynamic-range limitation discussed there. The mechanism behind it, which the main text does not give, is the one place where the binary health state is visibly implicated. `cellROS_rad51` assigns any damaged mutant cell directly to the nonviable state. *rad51*Δ therefore produces no cells in the intermediate damaged state at any dose, and its entire dose response must travel through the healthy-cell count.

#### S3. Fitting the alamarBlue kinetics model

*Corresponds to Materials and Methods, "AlamarBlue metabolism Michaelis-Menten reversible kinetics model" and "Parameter fitting for the kinetic model" in the main text, where this section is cited.*

**What is fit, and to what.** The kinetic model takes as its input the population growth curve produced by an AMMPER simulation and returns the concentration fractions of the three alamarBlue species over time. The six rate constants $v_1, v_2, v_3, K_1, K_2, K_3$ and the weight $k$ on damaged cells are shared across all twelve conditions, that is, six radiation doses in each of two strains. They are shared because the redox chemistry of alamarBlue is a property of the dye rather than of the genotype. Only the inoculum offset $g_0$ differs between strains. Nine parameters therefore account for twelve measured curves, and the strain difference is carried entirely by the population state that AMMPER supplies.

**How the parameters were estimated.** Parameters were estimated by global search using differential evolution (`scipy.optimize.differential_evolution`), with each offset subsequently refined by exhaustive scan with the kinetics held fixed. The bounds are the same for all twelve conditions and the random seed is recorded with the fitted values, so the search can be repeated exactly. Every parameter value reported in this work comes from that search. The choice of optimizer is not itself under test, since the same objective could be minimized by other means. What the fit is asked to reproduce is set by the objective and by the sharing of parameters across conditions, not by the search strategy.

**Error metric.** All errors reported in this work are the mean absolute deviation between predicted and measured concentration fractions, $\sigma = \frac{1}{n}\sum_{i=1}^{n} |x_i - y_i|$, averaged over the blue and the pink series and over all measured timepoints. This is the definition given in the main text Materials and Methods. Averaging rather than summing makes the value comparable across conditions irrespective of how many timepoints each contributes. The same definition is used in the fitting objective, in the tables below, and in every figure, so that all reported numbers are on one scale. The colorless species is not measured optically and therefore does not enter the error.

**Goodness of fit.** The table below gives the mean absolute error for each condition. The kinetic constants are identical in every row; the two strains differ only in $g_0$.

| Dose | Wild type | *rad51*Δ |
|---|---|---|
| 0 Gy | 0.029 | 0.097 |
| 2.5 Gy | 0.030 | 0.091 |
| 5 Gy | 0.023 | 0.053 |
| 10 Gy | 0.027 | 0.064 |
| 20 Gy | 0.025 | 0.075 |
| 30 Gy | 0.048 | 0.050 |
| mean | **0.030** | **0.072** |

Across all twelve conditions the mean absolute error is 0.051. The fit is uniformly good in the wild type, where the largest single-condition error is 0.048 at 30 Gy. In *rad51*Δ the residual discrepancy is concentrated at the two lowest doses (0 and 2.5 Gy) rather than at the high doses, and it takes a specific form. The measured curves flatten toward dose-dependent plateaus in the second half of the time course, whereas the predicted curves continue to descend. A model whose only dose-dependent input is the healthy-cell count cannot produce such a plateau; this is the same limitation discussed under dynamic range in the main text.

**Fitted values.** The estimated rate constants are $v_1, v_2, v_3 = 3.22, 18.6, 16.0$ and $K_1, K_2, K_3 = 1.39 \times 10^4, 218, 303$, with $k = 0.034$.

**Optimizer comparison.** We refit the same joint objective with two additional global optimizers, `scipy.optimize.dual_annealing` and CMA-ES (via the `cma` package). Each was run to its own convergence criteria rather than to a matched evaluation budget, so that no optimizer could be made to look worse only by being stopped early. The per-dose fit errors of the three searches are given in Fig. S7 and their fitted curves at the highest dose in Fig. S9. We disabled dual_annealing's gradient-based local-search refinement, since the admissibility constraint below creates a flat penalty plateau that a finite-difference gradient handles poorly. A Friedman test across the twelve paired dose and strain conditions found no significant effect of optimizer identity on per-condition fit error ($\chi^2 = 5.17$, $p = 0.076$, $n = 12$). Holm-corrected pairwise Wilcoxon signed-rank tests are reported for completeness (Fig. S8C): differential evolution against dual_annealing, $p = 0.085$; differential evolution against CMA-ES, $p = 0.339$; dual_annealing against CMA-ES, $p = 0.048$. We do not treat the last of these as evidence of a real difference, since it did not follow a significant omnibus test and $n = 12$ is a small paired sample. To check robustness beyond a single search, we reran each optimizer across five random seeds. CMA-ES converged to essentially the same fit regardless of seed (mean absolute error $0.0498 \pm 0.0000018$) and gave the numerically lowest mean error of the three; differential evolution was similarly stable ($0.0512 \pm 0.0001$); dual_annealing was more seed-dependent ($0.0537 \pm 0.0010$). We report the differential evolution fit throughout the main text, both because it was the search used throughout the rest of this work and because no practically meaningful or statistically supported advantage was established.

**Ordering of the curves by dose.** Because Figure 2 of the main text plots each dose on its own axes, the differences between doses cannot be read off it directly. Figs. S1 and S2 replot the predictions against the measurements on matched axes, one strain each, so that the separation between doses can be compared with the measured separation directly. Taking the value of each curve at the timepoint of greatest separation across dose, the rank correlation with dose is $+0.66$ for the predicted wild type blue series against $+0.89$ measured. For the pink series it is $-0.66$ against $-0.94$. The predicted ordering is imperfect, as the measured one is. For *rad51*$\Delta$ the prediction is perfectly monotonic in dose ($\pm 1.00$ for blue and pink) whereas the measurements are not ($+0.66$ and $-0.60$). The direction of the response is thus reproduced in every case. What is not reproduced is the magnitude of the separation, which is compressed roughly four- to fivefold in the wild type and ten- to twelvefold in the mutant (insets, Figs. S1 and S2).

**Where the compressed dynamic range comes from.** The compression originates in the population trajectory that AMMPER supplies rather than in the kinetic model. The simulated healthy-cell count at the end of the run varies by only 5.5% of its mean across the 0–30 Gy range for the wild type, and 7.9% for *rad51*$\Delta$. Direct damage is confined to the immediate neighborhood of the ion tracks, so most cells are never intersected by one. A minority of irradiated runs, 23 of 288 and all at 2.5 or 5 Gy, sustain no damage at all (Fig. S3). Two implementation choices compound this. The fitted weight on damaged cells is small ($k = 0.034$), so cells in the intermediate state contribute little to the predicted signal. `cellROS_rad51` also assigns a damaged mutant cell directly to the nonviable state rather than to the intermediate one. *rad51*$\Delta$ therefore registers no damaged cells at any dose, and its entire dose response must travel through the healthy-cell count. A graded health state with an explicit per-cell metabolic rate is the change these results point to, and it is a change to the cell mechanics rather than to the dye chemistry.

**Inoculum offset.** AMMPER begins each simulation from a single cell, whereas the plate-reader assay begins from a dense inoculum. A model fit to the full simulated trajectory would therefore have to account for several generations of near-zero signal that the experiment never observes. We therefore introduce one parameter, $g_0$, giving the point on the simulated growth curve that corresponds to experimental $t = 0$. Time is integrated continuously rather than on discrete generation ticks so that $g_0$ need not be an integer. It is a property of the assay setup rather than of the dye chemistry, and it is fit once per strain and held fixed across all six doses of that strain. For the wild type, $g_0 = 8.81$ generations, corresponding to approximately 463 cells at $t = 0$. Refitting $g_0$ separately for each dose gives values between 8.24 and 9.11 generations, a range of less than one generation, which is why a single shared value is used. For *rad51*$\Delta$ the fitted value is 6.21 generations, so the mutant population behaves as a wild type population approximately 2.6 generations behind, which is consistent with the slower net expansion expected of a repair-deficient strain. Per-dose refitting is looser for the mutant (4.97 to 7.27 generations) than for the wild type, which is consistent with the residual misfit above being concentrated in that strain. We retain the single shared value there too, rather than spending five further parameters to absorb it.

**Why the two strains are fit together.** We also fit the wild type in isolation and transferred the kinetics to *rad51*$\Delta$ with $g_0$ refit. On these data the two routes agree closely: the wild-type-only fit gives 0.0302 on the wild type and 0.0722 on the mutant, against 0.0302 and 0.0716 for the joint fit, or 0.0512 against 0.0509 over all twelve conditions. The kinetics recovered from the wild type alone therefore do transfer to the mutant without adjustment, which is itself informative. The dye chemistry inferred from one strain describes the other, and the strain difference really is carried by the population state. We report the joint fit because it is the more constrained arrangement, in that it cannot buy agreement on one strain at the other's expense. It is also the more stringent test of the claim that one dye chemistry plus one offset per strain accounts for all twelve curves.

**Identifiability.** The half-saturation constants $K_2$ and $K_3$ are both large relative to the amount of the corresponding substrate present, which places the reversible pink-to-colorless pair in an effectively first-order regime where only the ratios $v_2/K_2$ and $v_3/K_3$ are constrained. Scaling either pair together leaves the fit essentially unchanged: the wild-type mean absolute error is 0.0302 at scale factors of 1, 2, 10 and 100 alike for $(v_2, K_2)$ and likewise for $(v_3, K_3)$, agreeing to four decimal places in both cases. The same scaling applied to $(v_1, K_1)$ gives 0.030, 0.038, 0.055 and 0.059, nearly doubling the error. The flat response for the second and third pairs reflects the information content of the data rather than the bounds of the parameter search. $K_2$ and $K_3$ are therefore not individually identifiable from the present measurements, and we do not interpret their absolute magnitudes. $K_1$ is constrained by these data. That the reversible pink-to-colorless step should be the less well determined half of the chemistry is expected, given that the colorless species is not measured at all.

**Normalization of the experimental series.** Plate-reader absorbances are converted to concentration fractions by normalizing against the total dye present at $t = 0$, so that the measured blue and pink fractions together with the unmeasured colorless remainder sum to unity. This places the measurements on the same footing as the model output, which by construction conserves dye and so has its three species summing to one at every timepoint. The behavior of the normalized series is consistent with the assay chemistry. The measured blue-plus-pink sum is at its maximum at $t = 0$ in every one of the twelve conditions and decreases monotonically thereafter. The implied colorless fraction rises from 0.03–0.04 at $t = 0$ to between 0.18 and 0.58 at 15 hours, depending on strain and dose. Two conditions (*rad51*$\Delta$ at 0 and 2.5 Gy) exceed unity at $t = 0$ by 0.3% and 0.08% respectively, which is measurement noise in the raw absorbances at the first timepoint; we report this rather than rescaling it away. No part of the discrepancy between prediction and measurement is therefore attributable to the two being scaled differently.

**Mass conservation, and why the colorless species is constrained rather than clamped.** The three species are integrated without clamping, so that total dye is conserved to numerical precision. Clamping each species at zero instead, which is the more obvious implementation, silently creates dye whenever the reversible pink-to-colorless term runs backwards. For parameter sets in which the initial pink-to-colorless rate is negative, the clamp fires on the colorless species and total dye grows by more than 250% across the window. The model then degenerates into a two-species blue-to-pink conversion with the colorless species pinned near zero. This matters for interpretation as much as for arithmetic, because the assay chemistry requires the third species; a fit obtained under a clamp is not a fit of the model being described.

We therefore impose two admissibility constraints, evaluated before the error: a parameter set is rejected if the colorless species goes negative at any timepoint, and rejected if the colorless species never rises above 0.02 in concentration fraction over the window. The second constraint is necessary because the first alone is satisfied trivially by any parameter set that produces no colorless species at all.

We record that the search evaded weakly stated versions of these constraints on three separate occasions. In each case, it did so by driving parameters to a bound rather than by finding a better fit. The first evasion made the onward reduction rate negligibly small, so that the colorless species stayed just inside a tolerance. The second satisfied a growth requirement only in the final timestep. The third placed the inoculum offset at the end of the trajectory, where the population is saturated and the colorless species forms quickly enough to clear a threshold that is then met for the wrong reason. We mention this because it bears on how the reported errors should be read. Each of these produced a numerically improved objective while making the model less faithful to the chemistry, and each was found by inspecting the trajectories rather than the error values.

**Generation time.** Converting between simulated generations and experimental hours requires a generation time, which enters the model as a fixed constant rather than a fitted parameter. We use 198 minutes, the value implied by the duration of the 150 MeV proton experiments and the number of simulated generations. That is longer than the 80–120 minute doubling time typical of *S. cerevisiae* in rich medium at 30 °C. The discrepancy is expected, because the assay medium and temperature are not optimized for growth rate, and because the relevant quantity here is the net expansion rate of the population in the well rather than the minimum doubling time. The generation time is nonetheless a hyperparameter of the comparison rather than a measured property of the strain. It also trades off against $g_0$: a shorter generation time compresses the simulated trajectory onto the experimental window and is partly absorbed by a later offset. We have not attempted to fit the two jointly, since doing so would not be identifiable from these data.

**Availability.** The analysis code, the fitted parameters, the per-dose error tables reproduced above, and scripts regenerating every figure are provided with the submission.

#### S4. Converting absorbance measurements to aB concentrations

*Corresponds to Introduction, "Cell metabolism and alamarBlue (aB)" in the main text, where this section is cited.*

The BioSensor and the plate reader both report absorbances, whereas the kinetic model works in concentration fractions of the three alamarBlue species. Two corrections are needed to move between them. First, the absorbance spectra of resazurin (blue) and resorufin (pink) overlap, so the reading at either wavelength contains a contribution from the other species. Second, the wavelengths are not the same in the two settings. The BioSensor detects the oxidized and reduced forms with LEDs at 630 nm and 570 nm respectively, and tracks growth by optical density at 850 nm. The plate-reader measurements we fit to were taken at the intrinsic absorbance maxima of the two species, 570 nm for resorufin and ≈600 nm for resazurin, with growth read at 750 nm. The expressions below are written for the plate-reader wavelengths, since those are the data the model was fitted to. They therefore carry a ≈600 nm term corresponding to the resazurin maximum rather than to the flight LED, which is centered slightly off peak. They also carry a 750 nm rather than an 850 nm turbidity term. Applying the model to flight data would require the same expressions re-derived at the BioSensor wavelengths, with overlap coefficients measured there.

In the plate-reader data the turbidity is read at 750 nm, where neither dye species absorbs appreciably, and that reading supplies the cell-scattering background to be removed from the two dye channels. Write $A_\lambda$ for the absorbance at wavelength $\lambda$, and $\rho_\lambda = c_\lambda A_{750}$ for the scattering contribution at $\lambda$ implied by it. The coefficients $c_{600} = 1.04$ and $c_{570} = 1.06$ give the wavelength dependence of scattering over this narrow range. With $[B]$ and $[P]$ for the concentrations of the blue and pink species, the two species are recovered as

$$[B] = \frac{A_{600} - \rho_{600} - \alpha(A_{570} - \rho_{570})}{1 - \alpha\beta} \tag{S4}$$

$$[P] = A_{570} - \rho_{570} - \beta[B], \tag{S5}$$

where the constants $\alpha = 0.06$ and $\beta = 0.7$ are the pink:red and blue:green absorbance ratios, and account for the spectral overlap. The same 750 nm channel is the growth readout used to calibrate the gamma hit rate in Text S2. The resulting concentrations are then normalized against the total dye present at $t = 0$, which puts them on the same footing as the model output. That normalization and the checks on it are described under "Normalization of the experimental series" in Text S3.

#### Supplementary figures

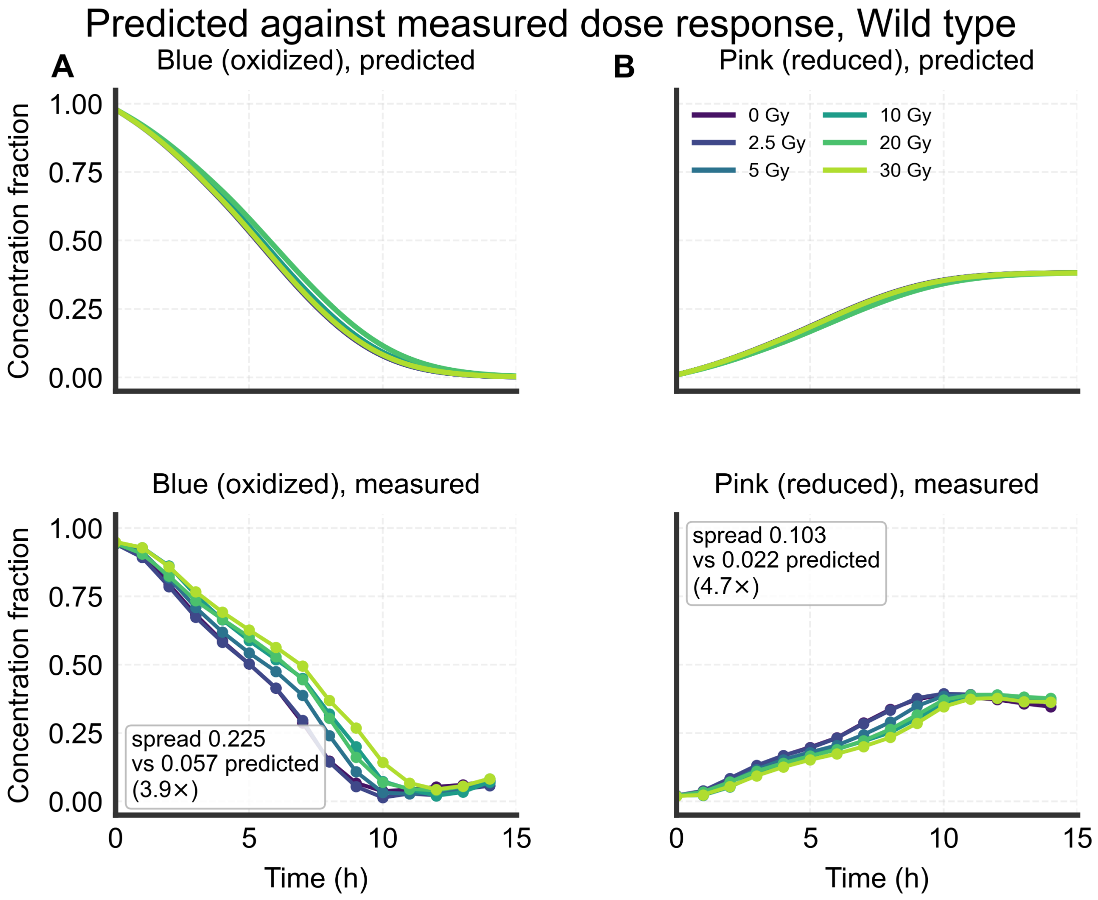


Figure S1: *Corresponds to Results, "AMMPER generates alamarBlue reduction curves using a naive Michaelis-Menten bulk model" in the main text, where this figure is cited.* **Wild type.** Dose dependence of the predicted and measured aB response, with predictions (top row) and measurements (bottom row) on matched axes so that the two can be compared directly without rescaling. Dose is indicated by color; the left column shows the blue (oxidized) species and the right column the pink (reduced) species. Insets give the measured and predicted peak-to-peak spread across dose at the timepoint of greatest separation, and their ratio. The predicted and measured curves are ordered by dose in the same direction in both species, but the predicted separation is four- to fivefold smaller than the measured one (0.057 against 0.225 concentration-fraction units for the blue species, 0.022 against 0.103 for the pink). The compression originates in the narrow spread of the simulated population trajectories across dose rather than in the kinetic model, which is quantified in Fig. S3 and discussed in Supplementary Text S3. The corresponding figure for *rad51*Δ is Fig. S2.

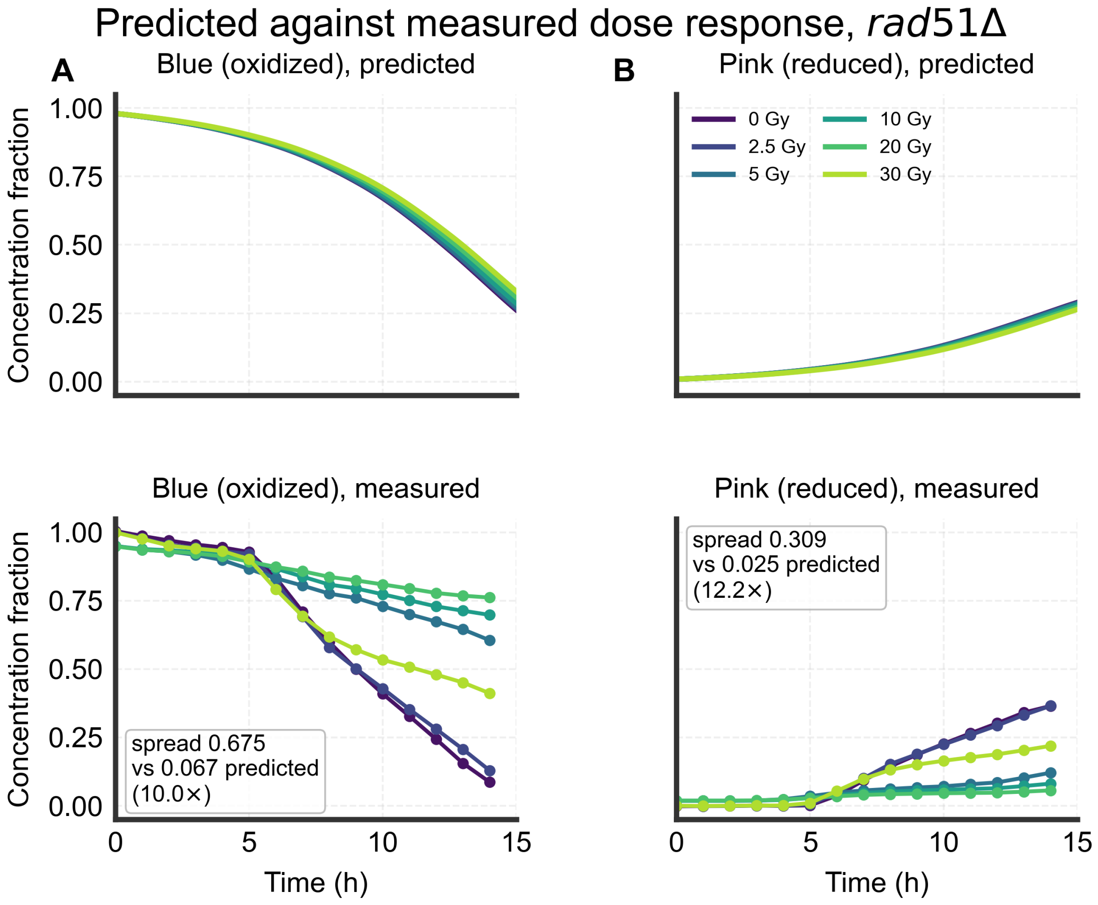


Figure S2: *Corresponds to Results, "AMMPER generates alamarBlue reduction curves using a naive Michaelis-Menten bulk model" in the main text, where this figure is cited.* ***rad51*** **Δ.** Dose dependence of the predicted and measured aB response, with predictions (top row) and measurements (bottom row) on matched axes so that the two can be compared directly without rescaling. Dose is indicated by color; the left column shows the blue (oxidized) species and the right column the pink (reduced) species. Insets give the measured and predicted peak-to-peak spread across dose at the timepoint of greatest separation, and their ratio. The predicted ordering is perfectly monotonic in dose (rank correlation ±1.00) whereas the measured curves are not, and the compression is correspondingly more severe: approximately tenfold for the blue species (0.067 against 0.675) and twelvefold for the pink (0.025 against 0.309). The compression originates in the narrow spread of the simulated population trajectories across dose rather than in the kinetic model, which is quantified in Fig. S3 and discussed in Supplementary Text S3. The corresponding figure for the wild type is Fig. S1.

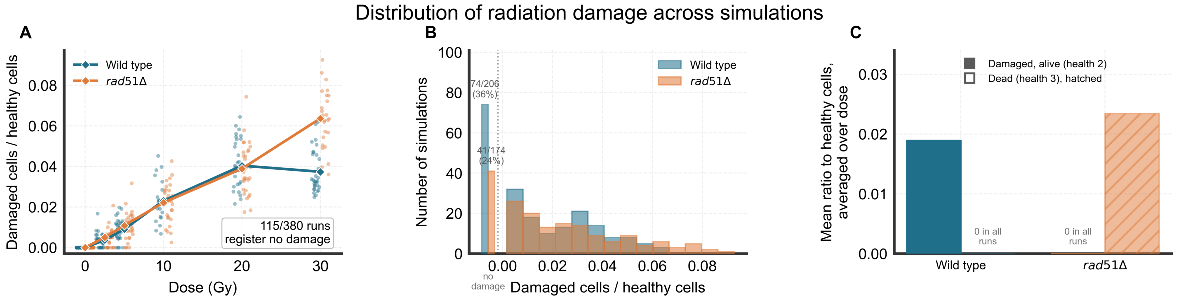


Figure S3: *Corresponds to Results, "AMMPER generates alamarBlue reduction curves using a naive Michaelis-Menten bulk model" and "Variance among AMMPER simulations is associated with localization of damage from ion tracks", where this figure is cited.* Distribution of radiation damage across all 380 archived proton simulations. **(A)** Per-run ratio of damaged to healthy cells at the end of the run, against dose, with every run plotted (horizontal jitter and a per-strain offset only; the vertical values are unperturbed) and the per-dose mean overlaid. 115 of the 380 runs register no damage at all. 92 of those are the two unirradiated 0 Gy conditions, which record no damage by construction; the remaining 23 are irradiated runs, 8 of 32 and 4 of 30 in the wild type and 8 of 29 and 3 of 30 in *rad51*Δ at 2.5 and 5 Gy respectively, in which the ion track (placed randomly at generation 2, when the population is four cells) missed the population. No run at 10 Gy or above, in either strain, is free of damage. The distribution is therefore zero-inflated and strongly non-normal, which is the justification for the rank-based tests used in the main text. **(B)** The same runs as a histogram of the per-run damage ratio, pooled over dose, one series per strain. The runs with no damage at all are drawn as separate bars to the left of the dotted break, labeled with their counts (74 of 206 wild type runs, 41 of 174 in *rad51*Δ), so that they are not pooled with runs carrying small nonzero ratios; the bars to the right of the break are the nonzero runs only. The distribution is therefore zero-inflated, with a short right tail and no central mode, which is why the analyses reported in the main text use rank-based rather than parametric tests. **(C)** The two damage states separately, averaged over dose. Each strain has exactly one nonzero bar, and the state that is empty in every run of that strain is annotated as such rather than drawn as a zero-height bar: the wild type accumulates cells in the damaged-but-alive state (health 2) and never a dead cell in any of 206 runs, while *rad51*Δ accumulates dead cells (health 3) and never a damaged-but-alive one in any of 174. This is a property of the implementation rather than of the data: `cellROS_rad51` assigns any damaged mutant cell directly to the nonviable state, so the mutant has no repairable-damage state by construction. It is the binary health-state limitation identified in the Discussion, made quantitative. Even at the highest dose the damage ratio remains small (mean 0.037 for the wild type at 30 Gy and 0.064 for the mutant), which is the origin of the compressed dose separation in Fig. S1.

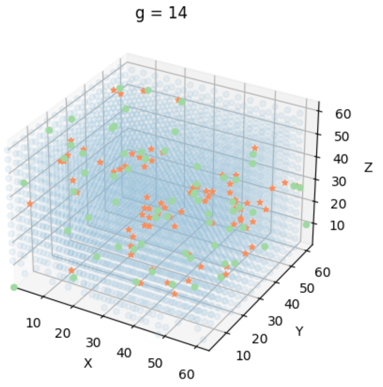


(a) Gamma radiation AMMPER visualization with *rad51*Δ cells.

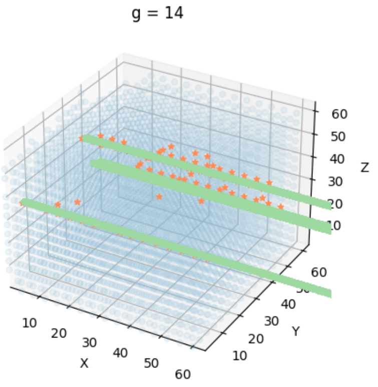


(b) Ion track AMMPER visualization, 150 MeV at 15 Gy, *rad51*Δ cells.

Figure S4: *Corresponds to Results, "The aB model transfers to a gamma radiation environment" in the main text, and to Supplementary Text S2, where this figure is cited.* AMMPER visualizations with *rad51*Δ cells. Healthy cells are depicted as blue circles, dead cells as red circles, and green circles or tracks represent the initial positions of ROS molecules generated by radiation events. **(a)** Gamma radiation causes non-localized damage to the cell population. **(b)** Ion tracks result in localized ROS generation along the tracks, with dead cells shown in red around the tracks. Healthy yeast cells, depicted in blue, are rarely affected if they are far from the tracks. This contrast in damage topology is what makes the gamma environment a distinct test of the model rather than a repeat of the proton one.

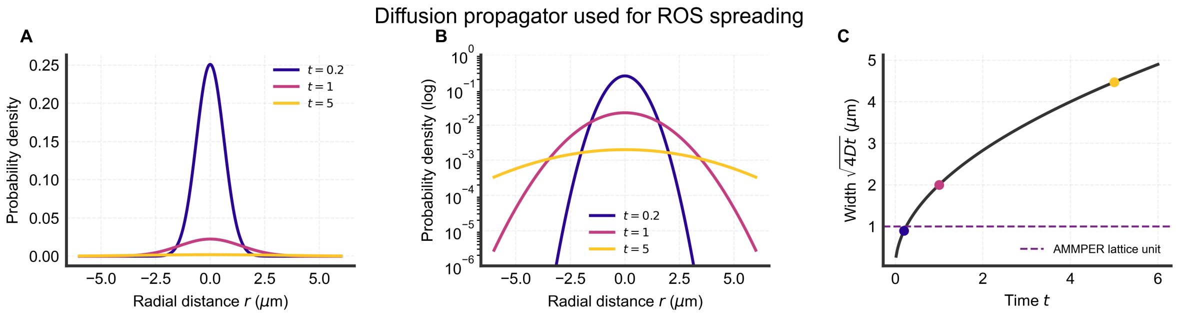


Figure S5: *Corresponds to Results, "Implementing decay and diffusion of reactive oxygen species produces little benefit over the naive model", and to Materials and Methods, "Diffusion and decay ROS molecule modeling", where this figure is cited.* Time evolution of the diffusion propagator (Equation S1), evaluated along a radial line for visualization at three times ($t = 0.2$, 1 and 5 in the arbitrary units of the figure, with $D = 1$). **(A)** Linear density axis. The probability density broadens as material initially localized at a point source disperses, and the peak height falls as $t^{-3/2}$, a factor of 125 across the times shown. **(B)** The same three profiles on a logarithmic density axis, which is where the broadening is legible: on the linear axis the $t = 5$ profile lies flat against zero. Both axes are shown because the linear panel is the faithful picture of relative magnitude and the logarithmic one the readable picture of shape. **(C)** The width $\sqrt{4Dt}$ implied by the propagator, with the three times marked and the AMMPER lattice spacing shown as a dashed line. Spreading narrower than one lattice unit cannot be represented on the discrete grid, which sets the shortest interval over which the diffusion model differs from the naive one.

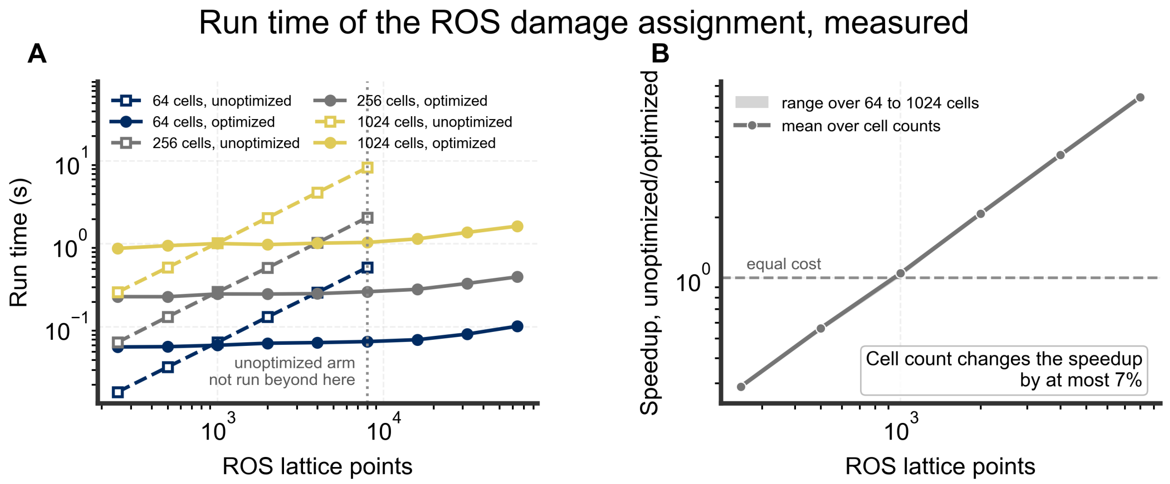


Figure S6: *Corresponds to Results, "AMMPER-2 requires reduced computational resources due to optimization of visualization process and ROS damaging mechanics", where this figure is cited.* Measured run time of the two ROS damage-assignment implementations, on identical inputs whose hit counts are asserted equal so that the comparison is of two implementations of one operation. "Unoptimized" is the interpreted nested loop over every ROS–cell pair used in AMMPER-1; "optimized" is the vectorized bounding-box filter used in AMMPER-2. Each point is the minimum of three repeats. **(A)** Run time against the number of ROS lattice points, for 64, 256 and 1024 cells. The unoptimized arm was not run beyond 8,000 points, marked by the dotted line, and the curve stops there rather than being extrapolated. **(B)** The ratio of the two, on the measured range. The vectorized implementation is slower below about 1,000 points, where a per-call filter overhead is not amortized, and faster above it by a factor that grows with problem size, reaching 8.0× at 8,000 points. This is the regime that matters in practice, since the diffusion ROS model emits 82 lattice points per deposition event where the naive model emits one, so enabling diffusion moves a simulation well to the right of the crossover. Measured on Linux 6.12.94 (x86_64), Python 3.12.13, NumPy 1.26.4, pandas 2.3.3.

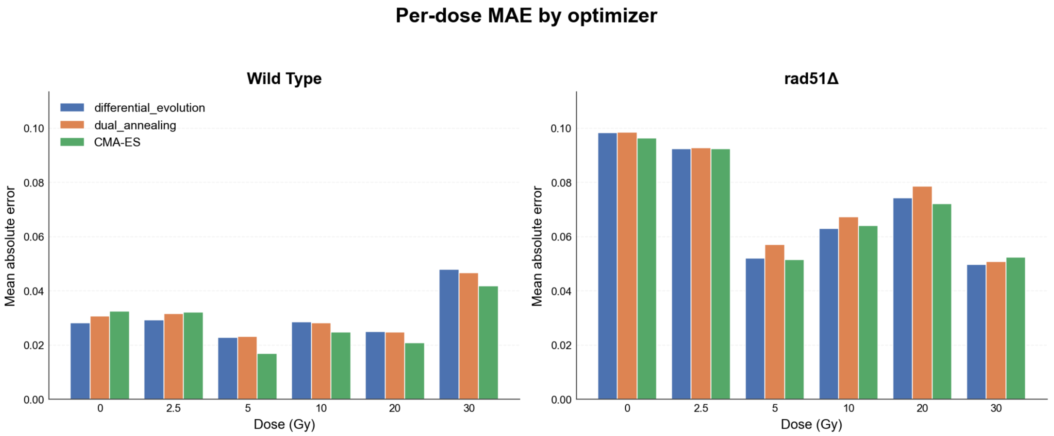


Figure S7: *Corresponds to Materials and Methods, "Parameter fitting for the kinetic model" in the main text, where this figure is cited.* Mean absolute error per dose for each of the three optimizers compared (differential evolution, dual_annealing, CMA-ES), shown separately for the wild type and *rad51*Δ. Summarized statistically in Fig. S8 and Supplementary Text S3.

**A. Mean absolute error per dose**

| | differential_evolution | | dual_annealing | | CMA-ES | |
|---|---|---|---|---|---|---|
| **Dose** | WT | rad51Δ | WT | rad51Δ | WT | rad51Δ |
| 0 Gy | 0.028 | 0.098 | 0.031 | 0.099 | 0.033 | 0.096 |
| 2.5 Gy | 0.029 | 0.092 | 0.032 | 0.093 | 0.032 | 0.092 |
| 5 Gy | 0.023 | 0.052 | 0.023 | 0.057 | 0.017 | 0.052 |
| 10 Gy | 0.029 | 0.063 | 0.028 | 0.067 | 0.025 | 0.064 |
| 20 Gy | 0.025 | 0.074 | 0.025 | 0.079 | 0.021 | 0.072 |
| 30 Gy | 0.048 | 0.050 | 0.047 | 0.051 | 0.042 | 0.052 |
| **mean** | **0.030** | **0.072** | **0.031** | **0.074** | **0.028** | **0.072** |

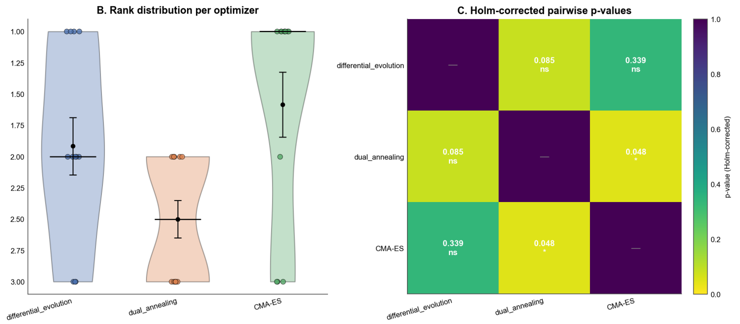


Figure S8: *Corresponds to Materials and Methods, "Parameter fitting for the kinetic model" in the main text, where this figure is cited.* **(A)** Mean absolute error per dose for each optimizer (differential evolution, dual_annealing, CMA-ES), wild type and *rad51*Δ shown side by side under each optimizer, with the mean per column given in the bottom row. **(B)** Per-condition rank distribution (1 = best fit) for each optimizer across the twelve dose strain conditions. **(C)** Holm-corrected pairwise Wilcoxon signed-rank $p$-values. A Friedman test found no significant effect of optimizer identity on fit error ($\chi^2 = 5.17$, $p = 0.076$); see Supplementary Text S3 for the full results, including a five-seed stability comparison.

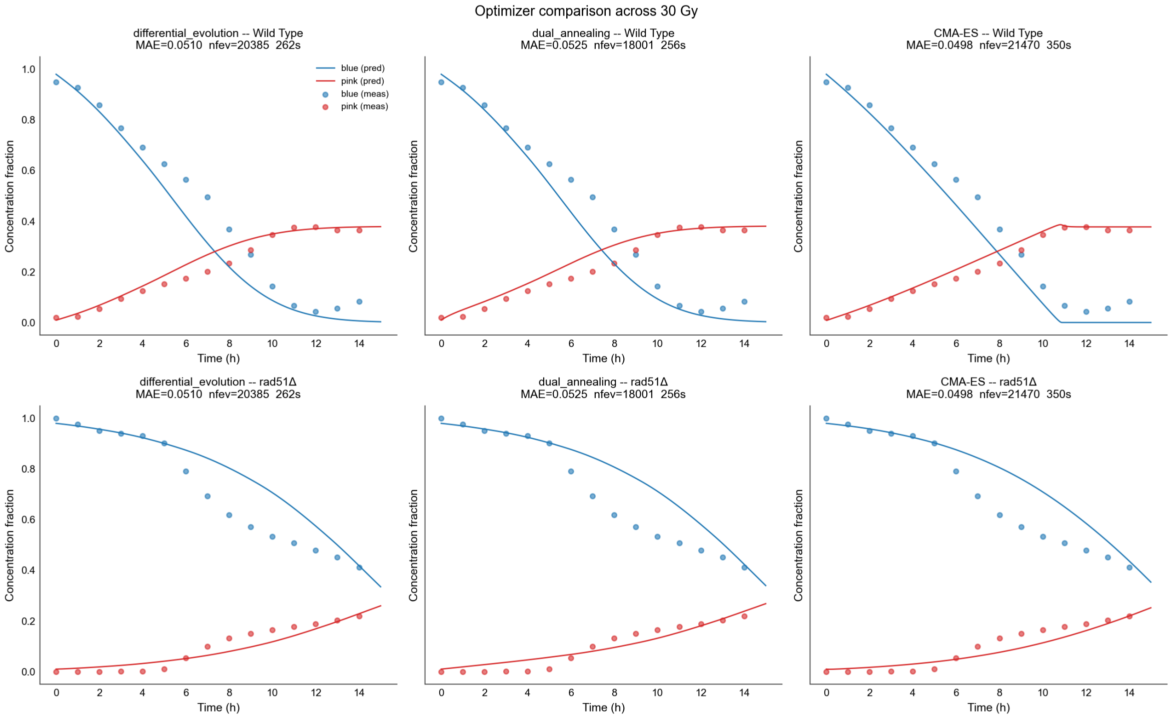


Figure S9: *Corresponds to Materials and Methods, "Parameter fitting for the kinetic model" in the main text, where this figure is cited.* Predicted versus measured aB response at the highest dose (30 Gy) for each strain, one column per optimizer, using each optimizer's own fitted parameters. Representative of the fit across all six doses; per-dose panels for every optimizer are not reproduced here but are archived in the repository, regenerated by `fit_ab_model_optimizer_comparison.py`.
